# A polyglutamine domain is required for *de novo* CIZ1 assembly formation at the inactive X chromosome

**DOI:** 10.1101/2020.11.10.376558

**Authors:** Sajad Sofi, Louisa Williamson, Gabrielle L. Turvey, Charlotte Scoynes, Claire Hirst, Jonathan Godwin, Neil Brockdorff, Justin Ainscough, Dawn Coverley

## Abstract

CIP1-interacting zinc finger protein 1 (CIZ1) forms large assemblies at the inactive X chromosome (Xi) in female fibroblasts in an *Xist* lncRNA-dependent manner. Here we address the requirements for assembly formation, and show that CIZ1 interacts directly with *Xist* via two independent domains in its N- and C-terminus. Interaction with *Xist* repeat E, assembly at Xi in cells, and the complexity of self-assemblies formed *in vitro,* are all modulated by alternatively-spliced exons that include two glutamine-rich prion-like domains (PLD1 and PLD2), both conditionally excluded from the N-terminal domain. Exclusion of PLD1 alone is sufficient to abrogate *de novo* establishment of new CIZ1 assemblies and Xi territories enriched for H3K27me3 in CIZ1-null fibroblasts. Together the data suggest that PLD1-driven CIZ1 assemblies form at Xi, are nucleated by interaction with *Xist* and amplified by multivalent interaction with RNA, so implicating a polyglutamine tract in the maintenance of epigenetic state.

## Introduction

X chromosome inactivation (XCI) is initiated by the long non-coding RNA (lncRNA) product of the X-linked gene *Xist* (X-inactive specific transcript) in the blastocyst of developing females (Brockdorff et al., 1992; Brown et al., 1992), leading to equalization of X-linked gene dosage between males and females (Penny et al., 1996). Once established gene silencing is maintained through subsequent cell generations, defining distinct initiation and maintenance phases of XCI (Wutz and Jaenisch, 2000). Initiation can be modeled in differentiating embryonic stem cells (ESCs), where recruitment of CIZ1 to the inactive X chromosome (Xi) is dependent on the repeat E region of *Xist* (Ridings-Figueroa et al., 2017; Sunwoo et al., 2017). Although this occurs concurrently with expression of *Xist* and with establishment of Xi chromatin, CIZ1 is not essential for XCI initiation and mice lacking CIZ1 develop normally (Ridings-Figueroa et al., 2017). CIZ1 becomes functionally relevant later, during maintenance of XCI.

In differentiated fibroblasts retention of *Xist* at Xi and maintenance of repressive chromatin modifications H2AK119ub and H3K27me3 (laid down by polycomb repressive complexes (PRC) 1 and 2 respectively) are dependent on CIZ1. At this stage CIZ1 forms large assemblies at Xi in female cells (Ridings-Figueroa et al., 2017) as well as much smaller nucleus-wide foci in both sexes (Ainscough et al., 2007). Deletion of CIZ1 has revealed a role in high-fidelity maintenance of PRC 1/2 gene sets, that is linked with a replication-coupled process of chromatin relocation (Stewart et al., 2019). At this point in the cell cycle CIZ1-Xi assemblies undergo a shift in properties that change its reliance on RNA for assembly integrity. This makes formation and stabilization of CIZ1-Xi assemblies of particular interest. Notably, loss of CIZ1 affects expression of approximately 2% of genes both X-linked and elsewhere in the genome (Ridings-Figueroa et al., 2017), suggesting that the mechanism by which it contributes to the preservation of the epigenetic landscape at Xi may be applicable to other CIZ1 foci and other loci.

It was recently hypothesised that *Xist*-dependent protein assemblies are phase-separated condensates that form a membrane-less compartment in the vicinity of Xi (Cerase et al., 2019). Membrane-less compartments, such as Cajal bodies and nuclear speckles, are micron-sized assemblies of proteins or RNA-protein complexes formed by liquid-liquid phase separation (LLPS) (Shin and Brangwynne, 2017). Most are sphere-like but others such as the TIS granule network form mesh-like structures (Ma et al., 2020). In most cases LLPS involves RNA-binding proteins (RBPs) harbouring prion like domains (PLDs). PLDs are intrinsically disordered regions with low sequence complexity that contain repeats of polar amino-acids such as polyglutamine that favour weak protein-protein interactions (Maharana et al., 2018). They play pivotal roles in normal cell physiology, however sometimes their physiological state is perturbed leading to abnormal protein aggregation, or maturation to amyloid-like fibres associated with neurodegenerative disease (Da Cruz and Cleveland, 2011).

Here we address the requirements for CIZ1 assembly at Xi in differentiated cells, and show that an alternatively-spliced PLD is required for *de novo* formation of functional CIZ1 assemblies at Xi, accompanied by repressive chromatin modifications. The data support the idea that these are localised at Xi by direct interaction with *Xist* via at least two independent CIZ1 interaction interfaces, one with preference for *Xist* repeat E.

## Results

### Alternatively-spliced prion-like domains contribute to CIZ1 assembly at Xi

Mouse CIZ1 (Fig. 1A) encodes two functionally distinct and partially characterized domains that we previously referred to as N-terminal replication domain (within amino acid 1-536 of RefSeq.NP_082688.1) which promotes cyclin-dependent initiation of DNA replication *in vitro* (Copeland et al., 2015), and C-terminal nuclear matrix anchor domain (within 537-845) which supports association with non-chromatin nuclear structures (Ainscough et al., 2007). Antibodies directed against epitopes in either the N- or C-terminal domains detect large assemblies of endogenous CIZ1 in Xi territories of wild-type (WT) female fibroblasts (Fig. 1B), activated lymphocytes and differentiated ESCs (Ridings-Figueroa et al., 2017; Sunwoo et al., 2017), but not in cells derived from CIZ1 null mice (Fig. 1B). Formation of CIZ1 protein assemblies at Xi can be modelled in primary embryonic fibroblasts (PEFs), when ectopic murine full-length GFP-CIZ1 is expressed from an integrated inducible expression vector (Ridings-Figueroa et al., 2017). Similarly, after transient transfection into cultured WT cells, full length GFP-CIZ1 assemblies are observed in Xi territories enriched for H3K27me3, but with variable efficiency depending on cell type; evident in 67% of cycling female 3T3 cells after 24 hours (Fig. 1C, Supplemental Fig.1A, B).

**Fig. 1.**
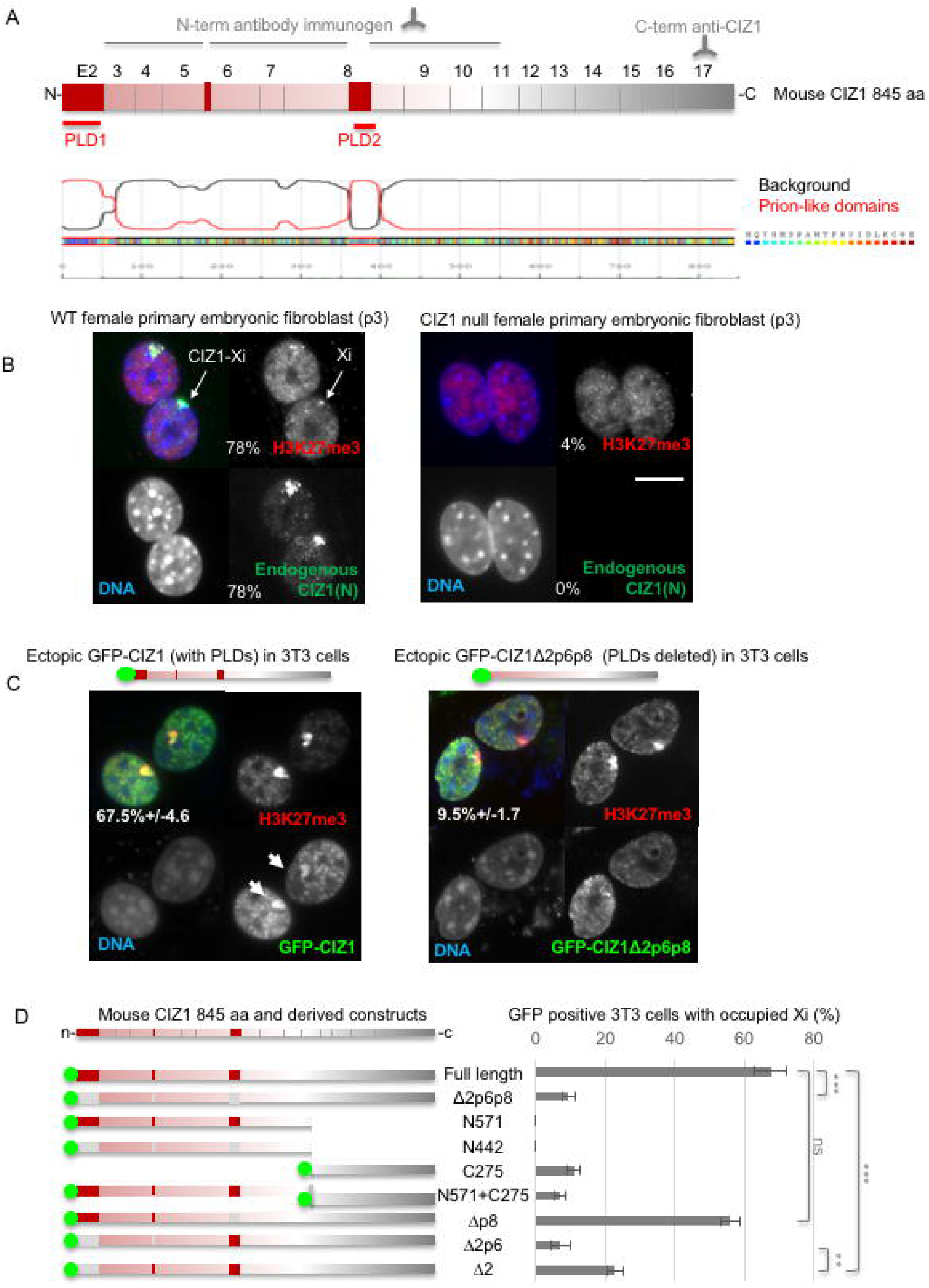
Alternative splicing of CIZ1 PLDs regulates assembly at Xi. A) Schematic showing translated exons 2-17, giving rise to the predicted full-length murine CIZ1, specified by RefSeq: NP_082688.1. Alternatively, spliced exon 2 excluded from CIZ1Δ2p6p8, as well as partially excluded exons 6 and 8 are indicated in red. The location of immunogens for anti-CIZ1 N-terminal domain antibody and anti-CIZ1 C-terminal domain antibody are shown above. Below, prion-like domains (PLDs) identified by PLAAC (Lancaster et al., 2014), align with alternatively spliced exons 2 (PLD1) and 8 (PLD2). B) Example images of murine primary embryonic fibroblasts (PEFs, passage 3) derived from WT and CIZ1 null mice (Ridings-Figueroa et al., 2017), showing endogenous CIZ1 detected with anti-CIZ1(N) antibody (green), H3K27me3 (red), and DNA (blue) in merged images. Bar is 10 microns. In this WT population of cycling cells 78% had colocalised CIZ1/H3K27me3-marked Xi’s, compare to 4% marked only with H3K27me3 in CIZ1 null cells (0% CIZ1). C) Expression of ectopic GFP-CIZ1 or GFP-CIZ1Δ2p6p8 (Coverley et al., 2005), 24 hours after transient transfection into WT cells. Arrows show accumulation of CIZ1, but not CIZ1Δ2p6p8, at sites of H3K27me3-enriched chromatin. The frequency with which CIZ1 assemblies are observed at Xi is indicated, with SEM. D) Left, illustration of CIZ1 deletion and truncation constructs missing combinations of exons 2, 6, 8, shown in red where present or in grey if deleted. Green circles, GFP. Right, their ability to assemble at Xi in cycling murine 3T3 cells, 24 hours after transfection, with SEM, and comparisons between key constructs by t-test. Example images for all constructs are given in Supplemental Fig. 1C, from the indicated number of repeat experiments (N).

In contrast, a naturally occurring alternatively-spliced variant of murine CIZ1 cloned from an embryonic day 11 cDNA library (previously termed embryonic CIZ1 or ECIZ1 (Coverley et al., 2005) and now designated CIZ1Δ2p6p8), is compromised in its ability to accumulate at Xi (Fig. 1C, Supplemental Fig.1A, B, C). Despite efficient nuclear targeting via a conserved nuclear localization signal (NLS) encoded by constitutive exon 7 (functionally validated in human CIZ1, Supplemental Fig. 2A, B, C), this variant does not form assemblies at H3K27me3-marked Xis with the same efficiency as CIZ1; evident in approximately 9% of cycling 3T3 cells after 24 hours (Fig. 1C and Supplemental Fig.1C). A similar comparison in WT female PEFs (p3) returned 62% and 30% respectively (Supplemental Fig.1A, B) confirming that alternative splicing modulates formation of CIZ1 assemblies at Xi.

CIZ1Δ2p6p8 lacks three sequence elements from its N-terminal domain (Fig. 1A), encoded by exon 2, and part of exons 6 and 8. Those encoded by exons 2 and 8 correspond to PLDs identified by *in silico* searches for prion-like amino acid composition (PLAAC) (Lancaster et al., 2014) (Fig. 1A), and are conserved in human CIZ1 (Supplemental Fig. 3A, B, C). PLD1 encoded by exon 2 is comprised of short (2-6 residue) polyglutamine (polyQ) repeats interspersed with leucine/isoleucine residues (Supplemental Fig. 3D), totaling 30 and 29 in human and mouse respectively. Expansion of CAG repeat elements, that encode glutamine residues, occurs in a group of genes that are linked with specific neurodegenerative conditions sometimes referred to as polyQ disorders. In unexpanded form these all normally encode a minimum of ten consecutive glutamines (Schaefer et al., 2012). Allowing for one mismatch within a run of at least 10 glutamines, Schaefer *et al* identified a more extensive set of proteins with polyQ tracts that represent 0.3% of the human proteome, enriched in nuclear functions. CIZ1 PLD1 fits this definition, identifying it as a polyQ tract. PLD2 is also enriched in glutamine residues but is not a polyQ tract. It is however subject to complex alternative splicing that results in at least three variants in humans (Supplemental Fig. 3A, Supplemental Table 1). Both PLDs are subject to conditional inclusion in CIZ1Δ2p6p8, and are therefore implicated in CIZ1 assembly at Xi.

### Role of PLD1 in CIZ1 assembly at Xi

To directly test which sequences are involved in CIZ1 assembly formation, we created a set of truncation and deletion constructs (Fig. 1D, Supplemental Fig. 1C), and screened them by transient transfection into wild-type female 3T3 cells which contain pre-existing endogenous CIZ1-XI assemblies. This showed them all to be expressed and present in the nucleus, and confirmed that N-terminal sequences alone, whether including (N571) or excluding (N442) the exons that are spliced out of CIZ1Δ2p6p8, are not sufficient for accumulation at Xi. The C-terminal anchor domain alone (C275) was observed to accumulate at existing Xis with low efficiency (Fig.1D), and not at all in cells that lack endogenous CIZ1 (see below). Co-transfection of N and C-terminal constructs (N571 and C275) did not reconstitute high efficiency targeting despite encoding the whole of CIZ1, indicating that both domains are required to be within the same polypeptide. Exclusion of exon 8 (PLD2) in the context of full-length CIZ1 had minimal effect (Fig. 1D), while deletion of exon 2 (PLD1) was sufficient to dramatically suppress assembly, an effect further augmented by co-exclusion of the five amino acids of exon 6. Similar analysis of the PLD1 deletion in WT PEFs (p3) confirmed its contribution to accumulation around H3K27me3-marked Xi chromatin (Supplemental Fig. 1A, B). This data highlights a central role for the polyglutamine tract that comprises PLD1 in formation of CIZ1 assemblies at Xi, but also shows that it is not sufficient.

### Initiation of new CIZ1 assemblies at Xi requires PLD1

Analysis in WT cells shows PLD1 to be involved in targeting to pre-existing CIZ1-Xi assemblies but what about *de novo* assembly in CIZ1 null cells? In primary fibroblasts lacking CIZ1, *Xist* is dispersed and both H3K27me3 and H2AK119ub are typically absent from Xi chromatin (Stewart et al., 2019). It should be noted that, during prolonged culture, both marks reemerge at Xi in CIZ1 null cells co-incident with upregulation of EZH2, so this analysis is carried out exclusively in early passage cells (p2-4) with a low (typically ~3% of cells) frequency of already-occupied Xi’s. It is against this baseline that change is measured during a 24-48 hour expression window. Re-expression of full-length CIZ1 from an inducible vector (Ridings-Figueroa et al., 2017), or by transient transfection (Fig. 2A) supports *de novo* assembly of typically one large CIZ1 aggregate per cell within 24 hours, accompanied by enrichment of H3K27me3 and H2AK119ub-marked chromatin (Fig. 2A), and retention of *Xist* within the assembly (Fig. 2B). Thus, CIZ1 is capable of establishing features of Xi chromatin in differentiated fibroblasts that are normally first established during embryogenesis. However, under the same conditions, GFP-CIZ1Δ2 is impaired in its ability to initiate new assemblies (present in 15% of cells that express it, compared to 40% for full-length CIZ1), and completely fails to support modification of Xi chromatin (3% enriched for H3K27me3 compared to 25% for full-length CIZ1, Fig. 2A). The data demonstrate that CIZ1 PLD1 is required for *Xist*-dependent repressive modification of Xi chromatin in differentiated fibroblasts.

**Fig. 2.**
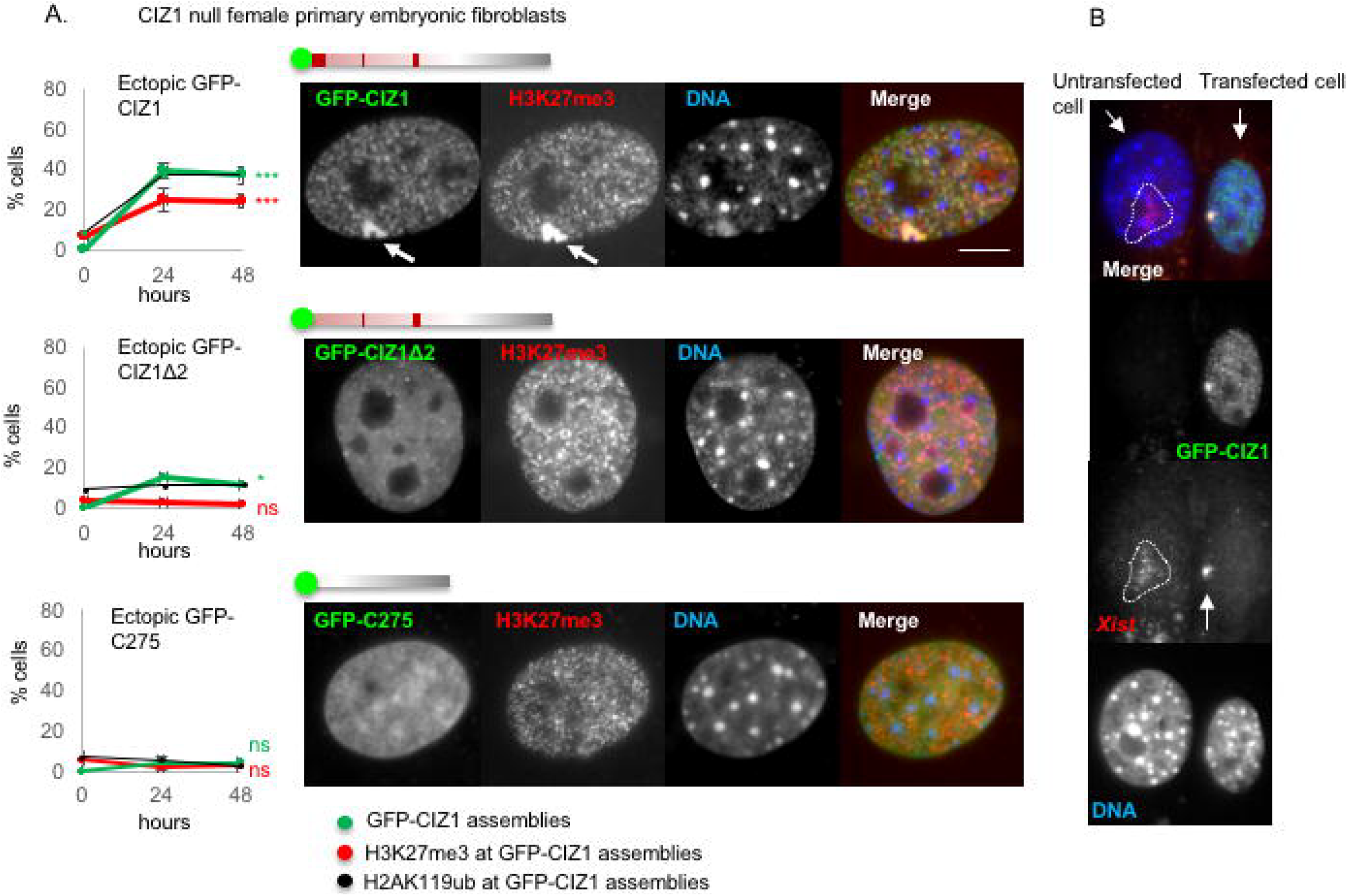
PLD1 is required to initiate formation of CIZ1-*Xist* aggregates in CIZ1 null cells. A) Frequency of GFP-CIZ1 (green) or H3K27me3 (red) marked Xis, 24 and 48 hours after transfection into CIZ1 null PEFs at p3. Absence of the polyglutamine rich PLD encoded by exon 2 from the CIZ1Δ2 construct (Δ1-68 in NP_082688.1) reduces accumulation at Xi and abolishes formation of H3K27me3-enriched Xi chromatin. Results are representative of 6 experiments with 3 independent isolates of primary cells (full-length CIZ1), and 3 experiments with 2 independent isolates of primary cells (CIZ1Δ2 and C275 constructs). Significance indicators refer to change over time for H3K27me3 frequency or CIZ1 assembly frequency. For full-length CIZ1 p<0.005 for CIZ1 assembly frequency at 48 hours, t-test. For H2AK119ub (black) results are from one experiment. Images show representative nuclei bearing GFP-fusion protein, co-stained for H3K27me3. B) *Xist* (red) in example cells with and without expression of GFP-CIZ1 (green), 24 hours after transient transfection, showing retention of *Xist* within *de novo* GFP-CIZ1 assemblies. Bar is 10 microns.

We also evaluated the GFP-C275 construct in CIZ1 null cells, and saw no evidence of accumulation at Xi or enrichment for H3K27me3 and H2AK119ub (Fig. 2A), indicating that its residual capacity to assemble at Xi observed in WT cells, requires the presence of endogenous CIZ1. Thus, the MATR3 domain (smart00451), and Jazz-type Zinc-finger (pfam12171) with predicted RNA-binding capacity, are not sufficient to support assembly. In fact, all of the sequence elements in C275, plus a further putative C2H2-type zinc finger (Supplemental Fig. 4B), are present in CIZ1Δ2, confirming that CIZ1’s zinc finger motifs are not sufficient to support functional assembly of CIZ1 at Xi in cells.

### CIZ1 binds Xist and prefers repeat E

*Xist* lncRNA is defined by a series of repeat motifs (A–F in the mouse, Fig. 3A) that interact with RBPs with functions in gene silencing (Monfort and Wutz, 2020; Nesterova et al., 2001). Our data and that of others indicate a functional relationship between CIZ1 and *Xist* repeat E (Ridings-Figueroa et al., 2017; Sunwoo et al., 2017), which consists of C/U/G-rich tandem repeats of 20–25 nucleotides long, over approximately 1.5 kb at the beginning of exon 7 of *Xist*. However, although high-resolution imaging shows that CIZ1 particles and *Xist* are in close proximity none of the existing data demonstrate a direct interaction. Moreover, the deletion studies of *Xist* (Ridings-Figueroa et al., 2017; Sunwoo et al., 2017), implicate different portions of repeat E in recruitment of CIZ1 to Xi. To test for a direct interaction, we generated *in vitro*-transcribed RNA probes to proximal (sense) and distal (sense and antisense) parts of *Xist* repeat E, as well as *Xist* repeat A for comparison (Supplemental Fig. 4A, Supplemental Table 2) and *Gapdh* and 18S ribosomal RNA (rRNA) as controls.

**Fig. 3.**
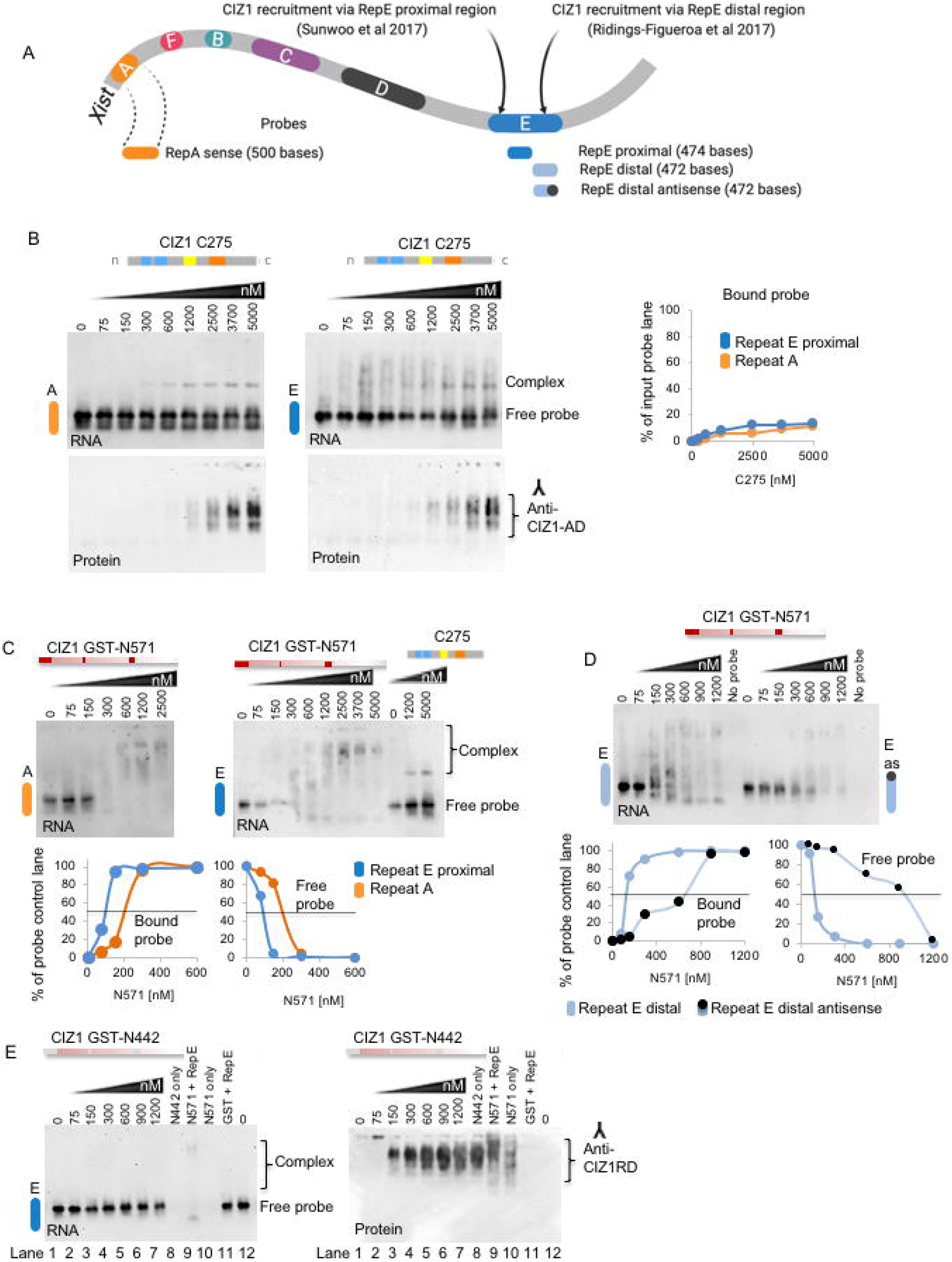
CIZ1 interacts directly with *Xist* repeat E. A) Schematic representation of domain organization in mouse *Xist* showing repeats A, F, B, C, D and E (Nesterova et al., 2001) and regions of repeat E thought to be involved in CIZ1 recruitment to Xi. Sense and antisense *in vitro*-transcribed RNA probes made from *Xist* repeat A and E regions are indicated below, showing proximal and distal E elements. B) Representative EMSAs showing binding of recombinant CIZ1 C-terminal fragment C275 with *Xist* repeat A or repeat E proximal region, and below western of the EMSA blot. Right, graph shows percentage of each RNA probe shifted by C275 at the indicated concentrations. C) Representative EMSAs showing binding of CIZ1 N-terminal fragment GST-N571, with repeat A and proximal repeat E (plus C275 shown for comparison). Below, graphs show percentage of bound and free RNA probe. D) Comparison of *Xist* repeat E distal sense and antisense RNA probes and their interaction with GST-N571. Below, graphs showing fraction of bound and free RNA. E) EMSA showing N-terminal fragment GST-N442 (lacking PLDs) and *Xist* repeat E proximal probe (plus GST-N571 and free GST controls). Interaction was detected with only GST-N571 control (Lane 9). Right, western blot of the EMSA showing GST-N571 alone (Lane 10) and in presence of repeat E proximal (Lane 9). Each EMSA is the representative of 2 or 3 independent experiments, and 0.3 nM of RNA probe was used in all experiments.

When analysed using electrophoretic mobility shift assays (EMSAs) we detected multiple discrete modes of binding between CIZ1 and RNA that were revealed by studying the N-terminal and C-terminal domains separately (Supplemental Fig. 4B). Recombinant C275 stably bound and shifted *Xist* repeat E or A to form a discrete RNA-protein complex, though with relatively low efficiency so that only 10-15% of input probe was complexed even under protein concentrations as high as 5 μM (Fig. 3B). We detected no difference in affinity between repeat E and repeat A probes, however C275 did not interact with *Gapdh* or *rRNA* probes (Supplemental Fig. 4C), indicating that its interaction with RNA is not promiscuous. CIZ1Δ2p6p8, which contains the whole C275 sequence, also formed a stable complex with *Xist* repeat E to a similar extent (Supplemental Fig. 4D). Thus, C275 can interact directly with *Xist*, but the data do not show preference for repeat E. To ask whether interaction with *Xist* is mediated by the Zinc finger domains in C275, we created a shorter construct encompassing just the C-terminal 181 amino-acids of CIZ1 (Supplemental Fig. 4B). Results of analysis by EMSA were indistinguishable from those with C275, arguing that the Zinc finger domains play no part in the observed interaction (Supplemental Fig. 4E, F. G).

Under the same conditions, the N-terminal 571 amino acids of full-length mouse CIZ1 (GST-N571) also formed stable complexes with both repeat E (proximal) and repeat A probes but with much higher affinity so that essentially all input probe was complexed and shifted in the low μM range (Fig. 3C, Supplemental Fig. 4H). Moreover, unlike C275, N571 showed heightened affinity for repeat E achieving half maximal shift at half the concentration of protein (100 nM) than was required for repeat A (200 nM). Similar results were achieved with the distal portion of repeat E, while anti-sense distal E showed lower affinity (Fig. 3D). GST-N571 also formed a complex with *Gapdh* to a similar extent as anti-sense distal E, but did not interact with *rRNA* (Supplemental Fig. 4I). Thus, GST-N571 interacts promiscuously with RNA but preferentially with *Xist* repeat E. In contrast, GST-N442 (lacking the PLDs) did not shift repeat E under the same conditions (Fig. 3E), showing that one of the direct interactions with *Xist* involves the alternatively-spliced domains in CIZ1.

Notably, GST-N571 did not form discrete RNA-protein species with any probe, and instead produced a broad smear indicative of a complex array of forms. The diffuse complexes most likely correspond to multiple nucleoprotein species, a conclusion arrived at by others (Grigoryev and McGowan, 2011). This could reflect protein-protein interaction, or multiple and variable numbers of CIZ1 molecules interacting with each RNA molecule, or a combination of both. Visualisation of input protein by western blot (Fig. 3E, right) shows that N571 exists as more than one species under native conditions (lane 10), and this is modulated by exposure to repeat E (lane 9).

Analysis of N and C-terminal domains in isolation from each other, therefore indicate at least two independent RNA interaction interfaces, both with some selectivity. No interaction was detected when GST was tested alone (Fig. 3E). Taken together these two-component interaction studies indicate direct multivalent interaction with *Xist*, and show that the N-terminal interaction has preference for *Xist* repeat E.

### CIZ1 forms condensates *in vivo* and self-assemblies *in vitro*

In the cellular context ectopic human GFP-CIZ1 initially forms nuclear foci similar to endogenous CIZ1, then coalesces into large droplets inside the nucleus over time, and ultimately kills host cells by a process that is not apoptosis (Higgins et al., 2012). Exclusion from the high RNA environment of the nucleus, via mutation of its NLS, influences droplet formation so that they form large aggregates in the cytoplasm (Supplemental Fig. 2C). Moreover, the N-terminal domain from murine CIZ1, with exons 2p6p8 (N571) and without (N442) exhibit a different propensity to coalesce, illustrated previously in male murine fibroblasts uncomplicated by events at Xi (Ainscough et al., 2007). Under these conditions both N-terminal domains coalesce as expression levels rise, but is first evident within one day for GFP-N571 compared to two days for GFP-N442. Thus, exons 2p6p8 influence the rate of droplet formation inside cells. In fact much of CIZ1 (not just its PLDs) is predicted to be structurally disordered (Supplemental Fig. 3C) a defining feature of proteins which undergo phase separation *in vivo* (Alberti, 2017; Shin and Brangwynne, 2017).

To explore and quantify these observed *in vivo* characteristics, we compared the properties of GST-N442 and GST-N571 *in vitr*o and recorded differences in their ability to condense and the nature of the structures formed. N571 formed irregular assemblies in a manner dependent on time and concentration (Fig. 4A, B, Supplemental Fig. 5A, B). Standardised quantification of phase contrast images after conversion to binary format (Fig. 4C) showed that N571 assemblies (at 10 μM) are first detectable at two hours, and continue to grow up to 48 hours. Particle number decreases as size increases, and this is accompanied by a decrease in circularity. Thus *in vitro*, GST-N571 undergoes spontaneous self-interaction independently of RNA to form microscopically visible assemblies of mean overall length of 6.2 microns (+/−0.6) at 24 hours.

**Fig. 4.**
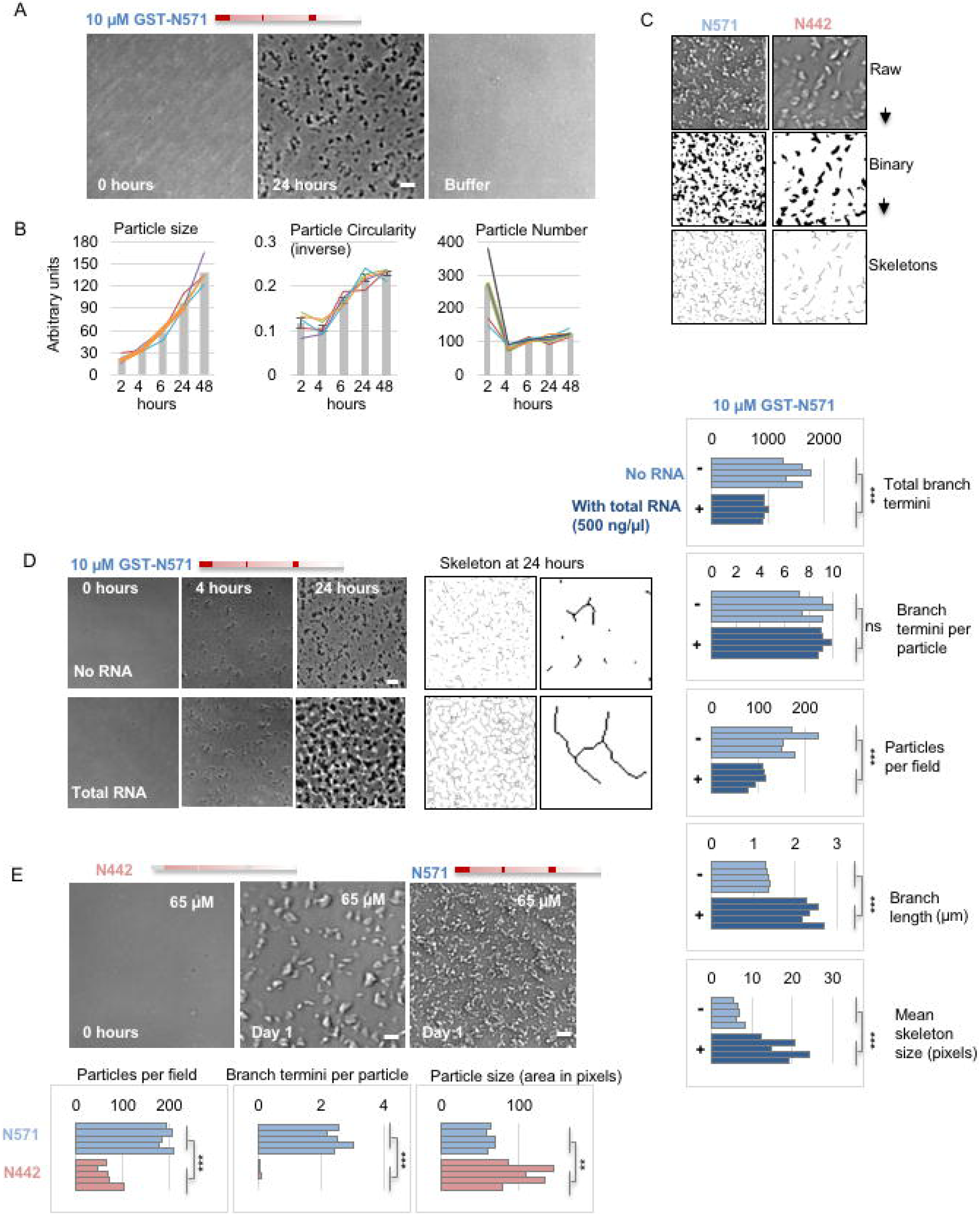
CIZ1 self-assembles into a mesh-like structure that is modulated by RNA. A) Phase contrast images showing CIZ1 N-terminal fragment N571 (10 μM) at the start and end of a 24 hour incubation.Bar is 10 microns. B) Quantification of particle size, number and circularity (expressed as inverse on a scale of 0-1) over time for samples of N571 at 10 μM. Histograms show mean of five samplings with SEM, and individual values as line graphs. C) Illustration showing conversion of phase contrast images (left N571, right N442) to binary format, and then to skeletons to derive quantitative information on ‘particles’ and ‘skeletons’ using FIJI (Schindelin et al., 2012). D) Images show N571 at 10 μM for the indicated times, with and without inclusion of total cellular RNA from murine 3T3 cells in incubations (500 ng/μl). Bar is 10 microns. Right, skeletonised images at 24 hours and close up illustrating branch structure. Histograms show quantification of five individual samplings with and without RNA, compared by t-test. E) Images of N-terminal fragment N442 (lacking PLDs) at 65 μM, before and after incubation. Bar is 10 microns. Comparison image of N571 at the same concentration is included. Below, graphs show results of five samplings showing differences in the number of particles formed, the degree of branching and the size of particles for N571 (blue) and N442 (pink).

Typically LLPS gives rise to spherical condensates, whereas CIZ1 assemblies resemble a fibrillar network. Skeletonisation of binary images (Fig. 4C) allows quantification of network complexity, returning information on the length and degree of branching. A direct comparison of N571 assemblies (10 μM at 24 hours) formed in the absence or presence of total cellular RNA (500 ng/μl) did not show suppression of assembly formation, and in fact lead to a quantifiable increase in skeleton ‘size’ (Fig 4D). This manifest as an increase in the maximum length of skeleton branches, a decrease in the overall number of particles and branch termini, but no effect on the number of branch termini per particle. Notably, inclusion of tRNA (500 ng/μl) rather than total RNA, did not drive an increase in branch length (Supplemental Fig. 5C). Thus, total cellular RNA promotes polymerisation of CIZ1, and possibly the formation of bridges between assemblies, but does not significantly increase their complexity. Neither RNA alone, nor BSA (in the presence or absence of RNA) formed microscopically visible assemblies under the same conditions (Supplemental Fig. 5D).

The behaviour of GST-N442 was different in several ways. A higher concentration (65 μM) was required to support assembly, and the structures that formed were larger, less complex and did not resemble a network, evident in branch scores (Fig. 4E). Thus, alternative splicing of CIZ1 influences assembly structure, shifting from branched interconnected assemblies for N571, to discrete entities for N442.

### Consequence of CIZ1 assembly at Xi

Proteomic and genetic screens have identified partially overlapping sets of proteins that interact with *Xist* inside cells, some of which have confirmed roles in *Xist*-mediated silencing (Chu et al., 2015; McHugh et al., 2015; Moindrot et al., 2015; Monfort et al., 2015). *Xist* interactors are enriched in glutamine-rich RBPs with a high probability of phase separating (Cerase et al., 2019), the potential of which has been hypothesized to play a role in the formation of the repressive Xi chromatin compartment. Supporting evidence emerged from analysis of the PTBP1, MATR3, TDP-43, CELF1 *Xist*-dependent protein assembly during ES cell differentiation. Its formation is driven by interaction with *Xist* repeat E during the later stages of initiation of X-inactivation, defining a time window when repeat E is essential (Pandya-Jones et al., 2020). This analysis also suggests that CIZ1 is recruited to Xi via repeat E independently of these factors, and we have shown that it is not required for the establishment of silencing (Ridings-Figueroa et al., 2017). An important question now is how the proteins that interact with repeat E relate to one another. Neither endogenous polypyrimidine tract-binding protein 1 (PTBP1) nor FLAG-matrin 3 (MATR3), both of which interact *in vivo* (Smola et al., 2016; Stork et al., 2019) (Pandya-Jones et al., 2020), were seen enriched around H3K27me3-marked Xis in fibroblasts (Supplemental Fig. 6). In fact, consistent with previous reports for MATR3 (Zeitz et al., 2009) not only do we see lack of enrichment, but also apparent exclusion from CIZ1 assembly zones (Supplemental Fig. 6C). Similar lack of enrichment was observed in primary embryonic fibroblasts for scaffold attachment factor A (SAF-A, also known as hnRNP U, Supplemental Fig. 6B), which interacts with *Xist* repeat D (Yamada et al., 2015). Lack of a positive relationship would argue against the suggestion that CIZ1 assemblies help concentrate other *Xist* interactors with a propensity to phase separate.

### Polyglutamine-mediated self-interaction

The role of CIZ1s polyglutamine domain in its assembly at Xi led us to question whether interfering with polyglutamine-mediated self-interaction would impact on characteristics of Xi implicated in maintenance of gene expression. Aberrant polyglutamine-mediated protein aggregation is well documented in relation to neurodegenerative disorders such as Huntington’s disease, where small molecule inhibitors have been trialled therapeutically. One such molecule, a cell-permeable amidosulfonamide compound, C2-8, inhibits polyglutamine aggregation *in vivo* when used in the micro molar (μM) range (Zhang et al., 2005). Consistent with a role for polyglutamine-mediated protein-protein interaction in CIZ1 assembly structure at Xi, C2-8 altered their shape (Fig. 5A, B) but not frequency (Fig. 5C). In treated cells, endogenous CIZ1-assemblies occupied a greater area and in some cells were evident as a flattened lamina-associated structure. An even more striking effect was seen with doxycycline-induced transgenic GFP-CIZ1 expressed against a CIZ1-null background (Ridings-Figueroa et al., 2017). Typically, this forms globular assemblies within 24 hours of induction, but their shape and frequency was affected by C2-8 in a dose dependent manner (Fig 5D). In the presence of C2-8, in some cells no GFP-CIZ1 assemblies were observed, or when present they were evident as an extended ribbon adjacent to the inner face of the nuclear lamina. In both endogenous and ectopic CIZ1 contexts we saw no evidence of reduced accumulation of H3K27me3 or H2AK119ub, which was evident in cells even with flattened CIZ1 assemblies (Fig. 5E), and we were still able to detect *Xist* in *de novo* CIZ1 assemblies after induction and treatment with C2-8 for 24 hours (Fig. 5F). While the effects of C2-8 on CIZ1 assembly structure in these experiments could be indirect, via interference with polyglutamine domains of other factors, it is nevertheless clear that polyglutamine-mediated interactions influence the shape and structure of the Xi chromatin territory. We speculate that those mediated by CIZ1 PLD1 could antagonize the reported interaction between *Xist* and lamin B receptor (LBR) (Chen et al., 2016) that normally anchors Xi at the nuclear periphery and might therefore influence the *Xist* and CIZ1-dependent transient internalization of Xi reported previously (Stewart et al., 2019; Zhang et al., 2007).

**Fig. 5.**
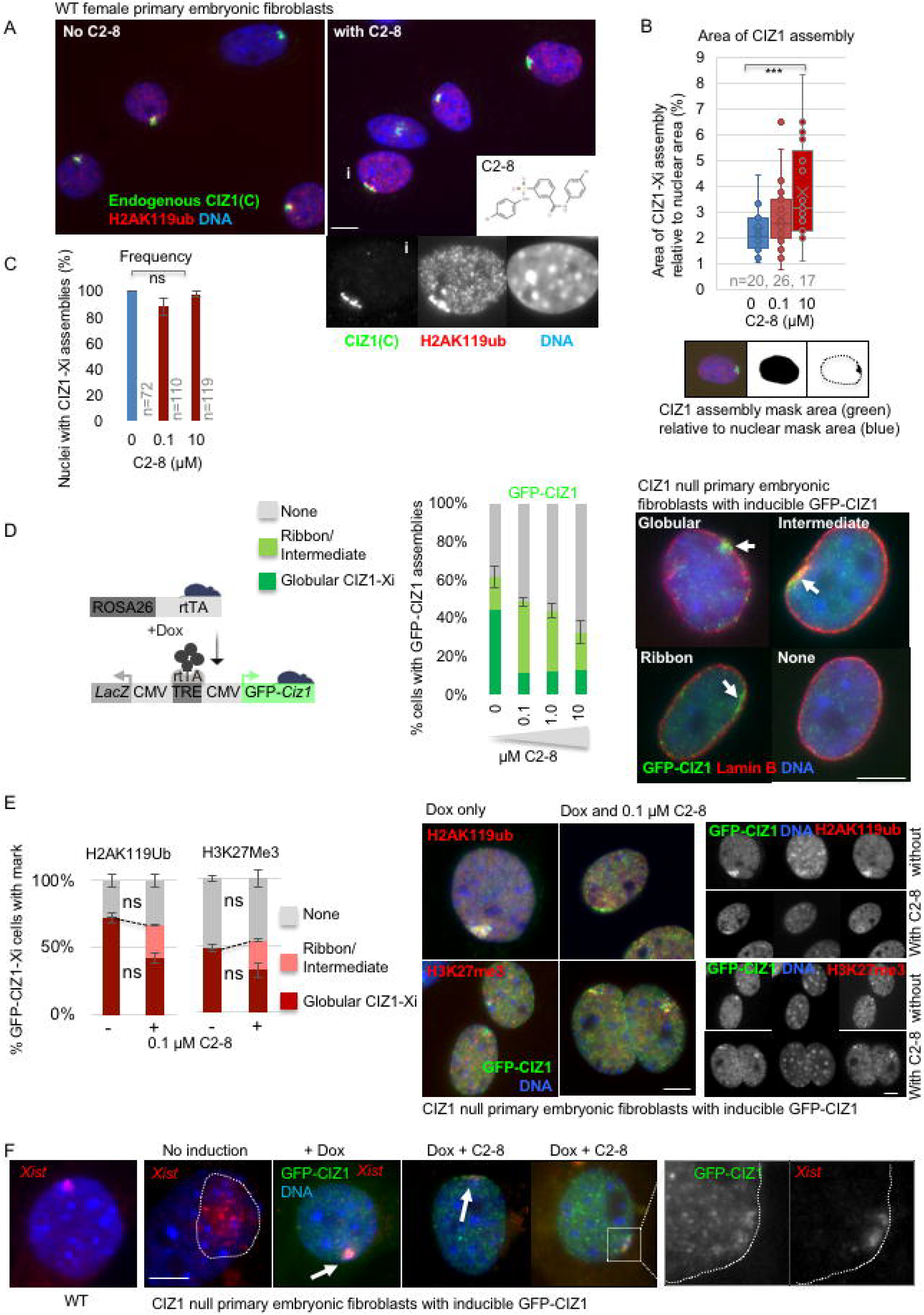
Polyglutamine-mediated interaction influences CIZ1 assembly structure at Xi. A) Example images showing endogenous CIZ1 (C-term, green) and H2AK119ub (red) in female WT PEFs at p3, without (left) and with (right) incubation with polyglutamine aggregation inhibitor C2-8 (inset) for 24 hours. DNA is blue. Bar is 5 microns. Below, example nucleus with i) lamina-associated ribbon-like CIZ1 assembly. B) Box and whisker plot showing area of CIZ1 assemblies (green signal), calculated as a proportion of nuclear area (blue signal), generated using image masks in FIJI (below). Data is representative of two experiments with independent WT primary cell isolates. C) Histogram showing that the frequency of CIZ1 assemblies remains unaffected by C2-8. D) Left, schematic of transgenes used to create doxycycline (dox)-inducible expression of full-length GFP-CIZ1 in CIZ1 null mice derived cells (Ridings-Figueroa et al., 2017). Tet-responsive element (TRE), CMV promoter (CMV), and reverse tetracycline transcriptional activator (rtTA). Histogram shows frequency of globular or ribbon-like GFP-CIZ1 assemblies (green), or absence of assemblies (grey) when cycling CIZ1 null PEFs at p3 were exposed to the indicated concentrations of C2-8 throughout a 48 hour induction period. Right, example images showing accumulation of GFP-CIZ1 (green) at Xi, either as a typical globular structure or an elongated ribbon-like structure associated with the nuclear lamina. Nuclei are counterstained for lamin B (red), DNA is blue, bar is 5 microns. Results are representative of three experiments. E) Histogram shows the proportion of GFP-CIZ1 assemblies that co-stain for H2AK119ub or H3K27me3 in female CIZ1 null PEFs (p3), in the presence or absence of C2-8, where ribbons are shown in pink and globular assemblies in red. Grey, no mark. Error bars are SEM from three replicates, ns, not significant, indicating that modification of Xi chromatin persists in assemblies whose structure is affected by C2-8. Comparisons are by t-test. Right, example images showing induced CIZ1-assemblies (green) and the indicated chromatin marks in red, DNA is blue. Bar is 5 microns. Far right, individual channels as indicated. F) Example images showing retention of *Xist* (red) upon induction of GFP-CIZ1 and the formation of *de novo* CIZ1 assemblies (green), in the absence and (two examples) presence of C2-8. Far right, high magnification, separate views of CIZ1 and *Xist* in grey scale. Far left, example WT cell showing normal *Xist* cloud.

## Discussion

The evidence presented here allows us to draw several conclusions about the behaviour of CIZ1 and to relate that to its function at Xi in differentiated murine fibroblasts. First, an alternatively spliced N-terminal polyglutamine PLD is required for formation of large sub-nuclear CIZ1-containing assemblies at Xi. In differentiated cells that were previously lacking CIZ1, this leads to accumulation of *Xist* and the acquisition of characteristic epigenetic marks. Second, CIZ1 can interact directly with *Xist* via at least two independent interfaces. One (C-terminal domain) is relatively low affinity, forms a single discrete complex and has no apparent preference for repeat E over repeat A (though does prefer *Xist* over *Gapdh*), while the other (N-terminal domain) is more efficient, prefers repeat E over repeat A, and is modulated by alternative splicing. Neither domain alone can support efficient accumulation at Xi inside cells, but together drive formation of CIZ1-enriched zones at Xi. Third, CIZ1 undergoes spontaneous self-association to form branched assemblies, also modulated by alternative splicing, and amplified by RNA. Together the data demonstrate the contribution of a polyglutamine domain to maintenance of X-inactivation, and potentially therefore in the stabilization of epigenetic state.

### Interaction with RNA

Some PLD proteins, such as Whi3, are capable of condensing on their own (Zhang et al., 2015) but multivalent interactions between PLD proteins and RNA typically modulate the properties of condensates, both by driving their formation and controlling morphology (Langdon et al., 2018; Maharana et al., 2018), while at high concentrations, such as those in the nucleus (Maharana et al., 2018), RNA can buffer against coalescence (Jain and Vale, 2017; Maharana et al., 2018; Yamazaki et al., 2018). Moreover, inside cells specific RNAs can seed intracellular assemblies controlling where condensation takes place (Zhang et al., 2015), for example paraspeckles form around the lncRNA *Neat1* (Hennig et al., 2015; Protter et al., 2018). For CIZ1, all three of these influences appear to be at play; i) exclusion from the nucleus is sufficient to drive droplet formation arguing that CIZ1 condensation may normally be buffered by the high RNA environment in the nucleus, ii) amplification and bridging of PLD-dependent *in vitro* assemblies is promoted by RNA, and iii) the location of *in vivo* CIZ1 assemblies is dependent on *Xist*, apparently mediated by direct interaction with *Xist*. The conclusion that alternatively spliced PLDs in CIZ1 contribute to a preference for repeat E is in line with recent studies which show that in RNA binding proteins with well-ordered RNA-binding domains, affinity for specific sequences can be driven by intrinsically disordered regions (Corley et al., 2020). This is also seen with DNA-binding specificity *in vivo* (Brodsky et al., 2020). Thus, our observations are entirely in line with emerging understanding of the contribution of disordered protein domains to protein-RNA interaction.

### Repeat E

A functional relationship between CIZ1 and *Xist* repeat E is evidenced by two studies that used *Xist* deletion analysis to implicate either its proximal (Sunwoo et al., 2017) or distal (Ridings-Figueroa et al., 2017) part. Here we independently tested two regions of repeat E and detected similar interaction with both *in vitro*, suggesting that any part may be sufficient to support interaction with CIZ1, but that not all of the constituent repeats are required. Despite a measurable preference for repeat E, the data also show that CIZ1 can interact, at higher concentrations, with repeat A, anti-sense E, and unrelated *Gapdh* RNA probes of similar length. Thus, the distinction is not strong *in vitro*, implying that relatively small differences in affinity for repeat E over other RNAs may be sufficient to capture CIZ1 *in vivo*. Alternatively, specificity may be achieved cooperatively with other factors *in vivo,* though there are currently no known interaction partners of CIZ1 that might confer such specificity. *In vitro* roughly half of RBPs, including PRC2 (Davidovich et al., 2015), bind RNA unspecifically (Jankowsky and Harris, 2015), and even the prion like RBP FUS, which nucleates around *Neat1,* exhibits unspecific RNA binding (Wang et al., 2015; Yamazaki et al., 2018).

It seems likely that much of the affinity for *Xist* repeat E reflects RNA structure-based determinants as we see no binding at all to highly ordered 18S rRNA (Anger et al., 2013; Rabl et al., 2011). In fact RNA structure has already been implicated in a polyQ-driven phase separation (Langdon et al., 2018; Zhang et al., 2015), and RNAs with large unstructured regions reported to form extensive intermolecular RNA-RNA interactions that play a role in formation of condensates with a mesh-like morphology (Ma et al., 2020). In the case of *Xist* its repeat E element is largely unstructured *in vitro*, and based on changes in SHAPE (selective 2′-hydroxyl acylation analyzed by primer extension) reactivity, appears to be a major protein-binding platform *in vivo* (Smola et al., 2016) that may seed nucleation of PLD-driven condensates.

### Model

Together our observations suggest a model in which multivalent interaction with RNA, coupled with PLD-driven self-association, support the formation of an RNA-CIZ1 matrix, localised to Xi by affinity for *Xist* repeat E (Fig. 6). We incorporate information on SAF-A which, like CIZ1, is reported to support retention of *Xist* at Xi. SAF-A interacts directly with AT-rich S/MAR DNA across the genome via its SAP domain (Gohring et al., 1997; Kipp et al., 2000), and with *Xist* via its RGG domain (Helbig and Fackelmayer, 2003), through which it is proposed to form a bridge between RNA and DNA that supports formation of the Xi territory during initiation of X inactivation (Hasegawa et al., 2010). This would remain unaffected by the presence or absence of CIZ1, but be insufficient to maintain enrichment of *Xist* at Xi in differentiated cells to the extent that an *Xist* cloud is detected by FISH. Indeed, *Xist* is spread across the nucleus in the absence of CIZ1 (Ridings-Figueroa et al., 2017), demonstrating that SAF-A alone has limited capacity to retain *Xist* within the Xi territory. Retention of additional molecules of *Xist* would be dependent on CIZ1, captured initially via affinity for *Xist* repeat E, but augmented by secondary interactions with other RNAs to form a CIZ1-RNA matrix.

**Fig. 6.**
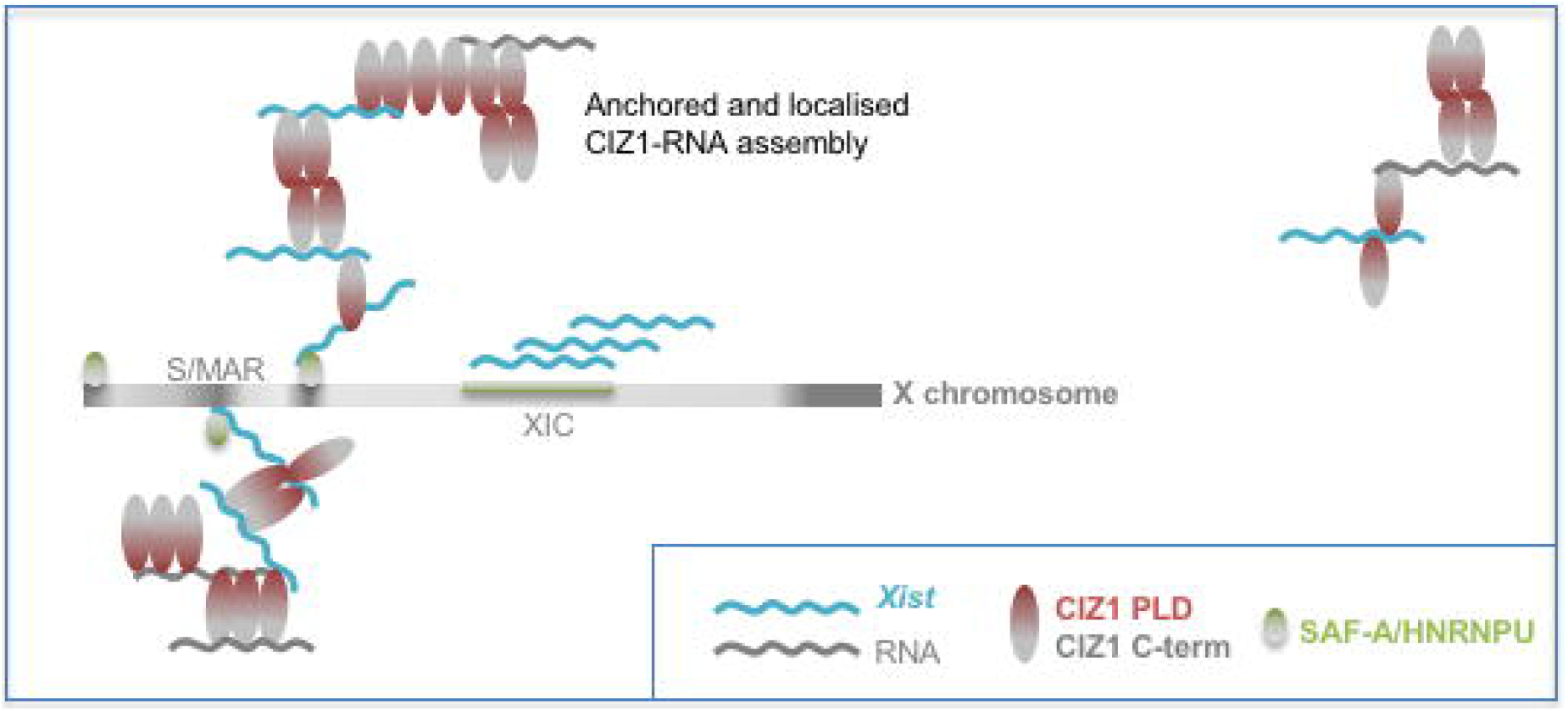
Model. Model illustrating the behaviour of CIZ1 at Xi, showing multivalent interaction with RNAs including *Xist* in the vicinity of Xi. SAF-A-anchored *Xist* at local S/MARS captures CIZ1 to form a protein-RNA matrix, that is amplified by unspecific interaction with other RNAs, and polymerization of CIZ1.

### Questions and limitations

The observations on the behaviour of CIZ1 that underpin this model pose some further questions. Notably, the same alternatively spliced PLDs support both self-association *in vitro* and high-affinity interaction with *Xist* in EMSA. What is the relationship between these events? For the purposes of our model we have assumed that both interactions take place at Xi, either independently or in the same molecule. However, from the available data it is not possible to assess whether the alternatively spliced domains all contribute to the same interaction interface or whether PLD1 interacts with *Xist* while PLD2 supports self-association, or vice versa. A related question is whether self-association precedes interaction with RNA, for example by creating a new RNA interaction interface, or indeed whether it is required. Moreover, we have not in this analysis addressed the possible contribution of the five amino acids of exon 6 that are missing from CIZ1Δ2p6p8. These do not meet the criteria of a PLD but do augment the Xi targeting defect caused by loss of PLD1.

More fundamentally, and despite clear evidence for dependence on PLD1, we cannot conclude that CIZ1-Xi assemblies form by LLPS or that they are fluid. Complicating this question is the observation that the stability (and possible fluidity) of CIZ1 assemblies fluctuates in the cell cycle, as revealed by temporally resolved sub-nuclear fractionation (Stewart et al., 2019). This shows that throughout most of the cell cycle CIZ1-Xi assemblies are independent of DNA because they remain unperturbed by removal of chromatin (identifying them as part of a classical ‘nuclear matrix’), but are dispersed upon digestion of RNA. Importantly, during a brief window in S phase, during or immediately after Xi chromatin replication, they become resistant to removal of RNA, as well as DNA, defining a transitional state that coincides with a shift in Xi location and maintenance of gene silencing (Stewart et al., 2019; Zhang et al., 2007). The molecular basis of this transient independence from RNA, and the regulation of CIZ1 assembly cohesion now deserve further attention.

Further questions also remain about the contribution and nature of the C-terminal *Xist* interaction. We show here that this is not mediated by CIZ1’s three zinc finger domains, but the precise determinants have not yet been delineated. Finally, while CIZ1Δ2p6p8 was originally cloned from an embryonic cDNA library, extensive CIZ1 splice variant diversity exists in both somatic and germ-line cells (Greaves et al., 2012), so we cannot yet say with certainty which forms are expressed during X-inactivation in the embryo.

### Implications for human disease

The data show that PLD1 is required for formation of new CIZ1 assemblies on Xi chromatin and for establishing repressive chromatin modifications, indicating a potential to impact on Xi gene expression. On the assumption that compromised assembly has similar consequences for maintenance of repressive chromatin as genetic deletion (Stewart et al., 2019), this suggests that CIZ1 alternative splicing, and indeed mis-splicing in human diseases, could have pleiotropic effects via compromised maintenance of epigenetic state. The first CIZ1 variant implicated in human disease, the paediatric central nervous system tumour medulloblastoma (Warder and Keherly, 2003), lacks exon 2 and thus PLD1. More recently, exome sequencing has revealed polymorphisms in the length of the polyglutamine tract in PLD1, with deletion of nine glutamines reported in seven malignant tumours of different origins. A range of other single nucleotide polymorphisms evident in both PLD1 and PLD2 are also listed in COSMIC (Tate et al., 2019) (Supplemental Table 1). Moreover, CIZ1-associated pathology is not limited to cancer. Sequence alterations that impact on PLD1, or which occur elsewhere and influence alternative splicing and tendency to aggregate inside cells, are also linked with familial cervical dystonia (Xiao et al., 2012; Xiao et al., 2014), and with Alzheimer’s disease (Dahmcke et al., 2008). Thus, modulation of CIZ1 nuclear assembly formation appears to play a role in diverse human pathologies, most of which have yet to be fully explored.

## Author Contributions

SS, DC, JA designed the experiments. SS, LW, CH, CS, JA, DC performed the experiments. JG, GT, NB contributed materials and advice. SS, DC wrote the paper. The authors declare no competing interests.

## Acknowledgments

We are grateful to Dr Han-Jou Chen and Dr Michael Plevin for critical comments on the manuscript, and to Matt Dowson for contributions to human cell analysis. This work was supported by a UK Medical Research Council grant MR/R008981/1.

## Methods

### Bacterial strains

All recombinant proteins were expressed and purified from the *E.coli* strain BL21-CodonPlus-RP. Cells were grown in Luria Broth (starter culture) and routinely cultured at 37°C while shaking at 220 RPM.

### Mouse cell lines

CIZ1 null mice were generated from C57BL/6 ES clone IST13830B6 (TIGM) harbouring a neomycin resistance gene trap inserted downstream of exon 1. The absence of *Ciz1*/CIZ1 in homozygous progeny was confirmed by qPCR, immunofluorescence and immunoblot. Mouse Primary Embryonic Fibroblasts (PEFs) were derived from day 13 or 14 embryos and genotyped essentially as described (Ridings-Figueroa et al., 2017). They were cultured in DMEM containing 10% foetal calf serum (PAAgold), 100 u/ml Penicillin, 10 μg/ml Streptomycin, 2 mM L-glutamine up to a maximum of passage 4. After passage 4 these cells are referred to as MEFs and were not used here. For inducible cells harbouring transactivator and responder transgenes, addition of doxycycline to media (10 μg/ml) was used to induce GFP-CIZ1, for 24-48 hours as indicated. Female 3T3 cells D001 were grown as described (Stewart et al., 2019) in DMEM (Sigma), 1% Penicillin, streptomycin, glutamine (Gibco),10% FBS.

### Expression constructs and transfection

Murine GFP-CIZ1 (845 amino-acids) and GFP-CIZ1Δ2p6p8 (formerly known as ECIZ1) in pEGFP-C3 (Clontech) were described previously (Coverley et al., 2005). Derived deletion constructs, N571 and N442 were generated by restriction digestion (Ainscough 2007). GFP CIZ1 C275 was made by ligating the 1 kb C-terminal XhoI fragment (Coverley et al., 2005) into the XhoI site of pEGFP-C2 (Clontech). GFP CIZ1 Δp8 (this study) was made by replacing the p8 containing BcuI/PmlI fragment of GFP CIZ1 845 with the Δp8 BcuI/PmlI fragment of GFP CIZ1Δ2p6p8. GFP CIZ1 Δ2p6 (this study) was made by replacing the Δp8 BcuI/PmlI fragment of GFP CIZ1Δ2p6p8 with the p8 containing BcuI/PmlI fragment of GFP CIZ1 845. GFP CIZ1 Δ2 (this study) was made by replacing the Δp6 KflI/PmlI fragment of GFP CIZ1 Δ2p6 with the p6 containing KflI/PmlI fragment of GFP CIZ1 845. The CIZ1 constructs, FLAG-MATR3 (Addgene 32880) (Salton et al., 2011), and FLAG-HNRNPU (Addgene 38068 (Baltz et al., 2012) were introduced into PEFs or 3T3 cells using Mirus X2 transfection reagent typically for 24 hours.

### Inhibitors

The amidosulfonamide inhibitor of polyglutamine aggregation C2-8 (N-(4-Bromophenyl)-3-[[(4-bromophenyl)amino]sulfonyl]benzamide, Sigma) was prepared at 10 mM in DMSO and used at the indicated concentrations in cell culture media for 24 or 48 hours as indicated.

### Protein expression and purification

Mouse CIZ1Δ2p6p8, C275, C181, N571 and N442 in frame with N-terminal tag glutathione S-transferase (GST) in pGEX-6P-3 expression plasmids (GE healthcare) were expressed in BL21-CodonPlus-RP *E. coli* using lactose-driven auto-induction. Cells were harvested after 24 hours at 20°C to produce 6-7 g of bacterial cell paste and frozen. Pellets were thawed on ice, suspended in 30-35 ml cold HEPES buffered saline (50 mM HEPES, pH 7.8 at 25°C, 135 mM NaCl, 3 mM EDTA, 1 mM DTT, supplemented with EDTA-free EZBlock™ protease inhibitor cocktail (BioVision) and 1 mM PMSF. Cells were sonicated on ice for 5 cycles (15 sec on, 30 sec off) at 60% amplitude using 6mm probe (microtip MS 73, Bandelin SONOPULS). Lysates were clarified by centrifugation for 20 minutes 4°C at 15000 rpm in Heraeus™ Multifuge™ X1 centrifuge with F15-6×100y fixed angle rotor. For affinity purification all steps were performed at 4°C. Clarified lysates were incubated with prewashed glutathione sepharose (GE Healthcare) and gently mixed with rotation for 1 h, then transferred to an equilibrated Poly-Prep® chromatography column, 0.8 × 4 cm (Bio-Rad). The column was washed with 10 column volumes (c.v.) of cold wash buffer 1 (50 mM HEPES, pH 7.8 at 25°C, 1 M NaCl, 3 mM EDTA, 1 mM DTT) followed by 3 washes with 10 c.v. cold wash buffer 2 (50 mM HEPES, pH 7.8 at 25°C, 135 mM NaCl, 3 mM EDTA, 1 mM DTT). Bead-bound protein was eluted with 4 × 2 c.v. of elution buffer (50 mM Tris-HCl pH 8.0 at 25°C, 10 mM L-glutathione reduced). Identity and purity of CIZ1-containing elution fractions was assessed by sodium dodecyl sulphate polyacrylamide gel electrophoresis (SDS-PAGE) with SimplyBlue™ safe stain (Invitrogen) and pre-stained protein ladder 10-250 kDa (Thermo Scientific). CIZ1-contaning fractions were pooled and reduced glutathione removed by buffer-exchange (50 mM Tris-HCl pH 7.0 at 25°C, 150 mM NaCl, 1 mM EDTA, 1 mM DTT) using Zeba™ Desalt spin columns (Thermo Scientific) following manufacturer’s instructions. GST tag was removed by incubating with 2 units PreScission protease (GE Healthcare) per 100 μg protein for 16-18 h at 4°C. Cleavage efficiency and specificity were examined by SDS-PAGE. Digestion mixture was passed over fresh glutathione sepharose and pure CIZ1 fractions were collected and concentrated with Vivaspin™ concentrator, 10-kDa cut-off (GE Healthcare) or Pierce® concentrator with 20-kDa cut-off (Thermo Scientific). Protein concentration was determined by absorbance at 280 nm with NanoDrop® ND-1000 spectrophotometer (Labtech, version V3.2.1). Absence of nucleic acids was verified by ensuring the ratio of UV absorbance at 260-280 nm was <0.7 and by visualization using denaturing gel electrophoresis. Immunoblot verification of purified CIZ1 proteins typically used 2 μg purified protein per lane, detected with antibodies listed in Supplemental Table 3. Purified proteins were supplemented to 9% v/v (final concentration) with sterile glycerol, aliquoted and snap frozen in nuclease-free cryo tubes (Nunc™) in liquid nitrogen and stored at −80°C.

### *In vitro* transcription of DIG labelled probes

DNA templates used for *in vitro* transcription of mouse *Xist* RNA were amplified from sequence verified plasmid pCMV-Xist-PA (Wutz and Jaenisch, 2000) (Addgene 26760), containing the murine *Xist* gene using high-fidelity Platinum® *pfx* DNA polymerase (Invitrogen™). PCR primers used for the amplifications contained T7 promoter sequences, were designed with SnapGene (GraphPad Software), and are shown in Supplemental Table 2. PCR amplicons of predicted size were confirmed by agarose gel electrophoresis, and purified using QIAquick® Gel Extraction Kit (Qiagen) for sequence verification. To generate human 18S rRNA probe, as an example of a highly structured template (Anger et al., 2013), the pTRI-RNA 18S control construct (MEGAshortscript™ T7 Transcription Kit, Ambion) was used as a DNA template for the *in vitro* transcription, to produce a 128 nucleotide product. To generate mouse *gapdh* RNA probe the *pTRI-GAPDH* control construct (NorthernMax™-Gly kit, Ambion) was used as a DNA template for *in vitro* transcription, to produce a 387 nucleotide product. *In vitro* transcription reactions were carried out with T7 RNA polymerase (MEGAshortscript™ T7 Transcription Kit, Ambion) at 37°C for 4 h. Upon completion, reactions were incubated with RNase-free TURBO DNase (Ambion) at 37°C for 15 min to digest template DNA, and RNA transcripts purified with MEGAclear™ Kit (Ambion) following manufacturer’s instructions. RNAs were eluted in nuclease-free water, 0.1 mM EDTA pH 8.0 and quantified by UV absorbance at 260 nm with NanoDrop® ND-1000 spectrophotometer (Labtech, version V3.2.1). Purity, transcript size and integrity of all RNA constructs were examined under denaturing conditions with 7M Urea-PAGE gels stained with SYBR™ Safe DNA Gel Stain (Invitrogen), with RNA Century™-Plus Markers 0.1–1 kb (Ambion). All *in vitro* transcribed RNA transcripts were DIG-labelled by incorporating digoxigenin-11-UTP at a ratio of 1:15 with unmodified UTP during transcription.

### Electrophoretic mobility shift assay (EMSA)

In a 10 μl binding reaction, purified recombinant CIZ1 proteins and derived fragments (CIZ1Δ2p6p8, C275, C181, N571, N442) were incubated with DIG-labelled RNA probes in binding buffer (10 mM Tris-HCl, pH 7.5 at 25°C, 30 mM NaCl, 2.5 mM MgCl_2_, 0.2 mM DTT, 0.1% IGEPAL® CA-630 (Fluka™), 0.1 mg/ml yeast tRNA (Ambion®), 0.4 units of RNaseOUT™ (Invitrogen™), 1% v/v glycerol) at 30°C for 20 min. Before use RNA was denatured at 80°C for 3 min and snap cooled on ice for 2-3 min to allow RNA refolding. Reaction mixtures were loaded onto an 11 cm × 6 cm horizontal nondenaturing 0.7% agarose gel (Molecular Biology Grade agarose, Melford) buffered with 1× filter sterilized TBE at 4°C. Gel electrophoresis was carried out for 60 min at 6 V/cm in an ice box in a 4°C cold room. Blotting was performed by upward capillary transfer onto positively charged nylon membrane (Hybond-N+, Amersham). Membrane and RNA were cross-linked by exposure to shortwave UV light at 254 nm at 120 mJ (GS Gene linker UV Chamber, BIO-RAD). Membranes were washed in 1× wash buffer (0.1 M maleic acid, 0.15 M NaCl, pH 7.5 at 20°C, 0.3% (v/v) Tween-20) for 5 min under agitation and next blocked in 1× blocking buffer (Roche) for 30 min at room temperature with gentle shaking. A polyclonal sheep anti-digoxigenin antibody conjugated to alkaline phosphatase (Roche) was added to membrane at 1:10,000 dilution then washed twice in 1× wash buffer at room temperature with gentle shaking and equilibrated in 1× detection buffer (100 mM Tris-HCl, pH 9.5, 100 mM NaCl) for 5 min at room temperature. The membrane was developed by adding chemiluminescent CSPD substrate (Roche) at 0.25 mM final concentration at room temperature for 5 min and incubated at 37°C for 10 min for alkaline phosphatase activation. The chemiluminescent signal was acquired with a PXi touch Chemiluminescence imaging system (Syngene). Densitometry was carried out with GeneTools software (Syngene, version 4.3.8.0). All quantifications were expressed relative to the lane containing RNA probe but no protein for each gel.

### *In vitro* assemblies

Purified proteins in 50 mM Tris-HCl, pH 7.0, 150 mM NaCl, 1 mM DTT were concentrated to greater than 60 μM stock solution, then diluted in isolation buffer as appropriate. Incubations were carried out in uncoated multi-well plates at room temperature, unless indicated otherwise. Total cellular RNA was isolated from a female 3T3 cell line (Ridings-Figueroa et al., 2017) using Trizol reagent as recommended. Transfer RNA was from yeast (Ambion®). Samples were imaged using an Evos Xl (AMG) light microscope fitted with an Evos fluorite LWD phase contrast 40× 0.65 objective, using constant illumination settings and capture times within a sample series to generate 2048×1536 9.4MB TIFF image files. Image sets were processed using FIJI to produce optimized contrast for reproduction purposes.

### RNA Fluorescence in situ hybridization

Cy3-labelled RNA FISH probe was produced from mouse pCMV-*Xist-*PA (Wutz and Jaenisch, 2000) and purified using GENECLEANⓇ kit (MPBIO). An 11 kb Spe1-Sal1 fragment (mid exon 1–7) was labelled with Cy3-dUTP (GE Healthcare UK Ltd) using Bioprime labelling kit (Invitrogen). 100 ng *Xist* fragment was added to 20 μl 2.5X random primer buffer and 9 μl nuclease free water, denatured by boiling for 5 minutes, then incubated on ice for 2 minutes. dNTP’s (5 μl dATP, dCTP and dGTP, 3 μl dTTP and 1 μl Cy3 labelled dUTP) were added under low light, then 1 μl Klenow fragment, and incubated overnight at 37°C in the dark. After incubation, 5 μl stop buffer, 10 μl Cot1 DNA (Invitrogen) and 5 μl salmon sperm DNA (Novagen) were added. Labelled DNA was precipitated twice with 3 M NaOAc and 100% ethanol to remove unincorporated nucleotides, resuspended in 80 μl hybridisation buffer (50% formamide, 10% Dextran sulfate, 2 mg/ml BSA, 2X SSC) and stored at −20°C. Cells were seeded onto glass coverslips at ~70% density and transfected as required. After 24 hours cells were washed in RNase free PBS and fixed in fresh 4% paraformaldehyde, on ice, for 15 minutes. Cells were washed in PBS (3 x 5 minutes), then incubated in 1 ml permeabilization solution (PBS+0.5%Triton X-100 (Sigma), 0.5% BSA (Jackson ImmunoResearch) with 10 mM VRC (NEB) per coverslip for 10 minutes at room temperature. The cells were washed in PBS (3 x 5 minutes) and stored at 4°C in 70% ethanol. Cy3-labelled probe (10 μl per coverslip) was defrosted on ice, and VRC added to 10 mM. The probe was denatured (74°C, 10 minutes), spun briefly and incubated at 37°C for 20 minutes. Prepared cells were dehydrated through an alcohol series of 70%, 95% and 100% ethanol for 2 minutes each. Coverslips were air dried for 5 minutes, placed cell side down onto 10 μl denatured probe on an RNase free slide, sealed with rubber cement and incubated at 37°C overnight in a humidified chamber in the dark.

Coverslips were carefully lifted and sequentially washed with 2X SSC, 50% formamide (3 × five minutes, 39°C), and 2X SSC (3 × five minutes, 39°C), then once each in 1X SSC and 4X SSC for 5 minutes at room temperature. All washes were supplemented with VRC. Following a brief dip in DEPC water, coverslips were mounted in Vectashield with DAPI, or processed for immuno-FISH.

### Immunofluorescence

For immunofluorescence, cells were grown on coverslips then washed in PBS and fixed in 4% PFA to reveal total protein, or alternatively briefly washed in PBS with 0.1% Triton-X 100 then fixed in 4% PFA, to reveal the immobilised protein fraction (‘detergent treated’). For combined RNA/immunoFISH cells were processed for RNA as described then continued as below. Coverslips were blocked in AB (1x PBS, 10 mg/ml BSA, 0.02% SDS, 0.1% Triton-X 100), for 30 minutes then incubated with primary antibodies for 2 hours, 37°C, washed in AB, incubated with secondary antibodies for 1hr, 37°C, washed and mounted on glass slides with Vectashield medium containing DAPI (Vector labs). All antibodies used are detailed in Supplemental Table 3. Fluorescence images were captured using a Zeiss Axiovert 200M fitted with a 63X/1.40 Plan-Apochromat objective and Zeiss filter sets 2, 10, 15 (G365 FT395 LP420, BP450-490 FT510 BP515-565, BP546/12 FT580 LP590).

### Ethics

All work with animal models is compliant with UK ethical regulations. Breeding and genetic modification of mice were carried out under UK Home Office license and with the approval of the Animal Welfare and Ethical Review Body at the University of Leeds and Oxford. Analysis on cells and tissues derived from these mice was carried out with the approval of the Animal Welfare and Ethical Review Body at the University of York.

### Bioinformatics

Protein motifs were identified using Psort II (https://psort.hgc.jp/). Alignments were performed using Clustal omega. Protein disorder was predicted using PONDR (http://www.pondr.com/) and disEMBL (http://dis.embl.de/), and prion-like domain using PLAAC (http://plaac.wi.mit.edu/).

### Densitometry analysis

For EMSA experiments, densitometry was performed with Gene Tools software and quantifications were expressed relative to the lane containing only RNA probe. The percentage of shifted RNA and number of experiments are given in legend of Figure 2.

### Quantification and analysis of *in vitro* assemblies

Samples were imaged using Evos Xl (AMG) light microscope. Images were processed with Fiji to perform binary and skeleton analysis of protein particles. Quantitative analysis was performed on non-adjusted images as follows. For ‘particle size’, number and circularity measurements, 250×250 pixel image sections were converted to binary output using FIJI and all particles (0-infinity) analysed. Circularity reflects the smoothness of the perimeter of an object, where a perfect circle has a circularity value of 1.0, using the formula 4pi(area/perimeter^2). For branch analysis, binary images were converted to skeletons and analysed without pruning. At least 5 images were analysed for each condition and results summarized to yield mean data for each parameter, including branch termini (end-point voxels), and branch length (max branch length per particle expressed as average per field), or non-junction non-terminus skeleton body (slab voxels), referred to as ‘skeleton size’. Note, that highly assembled particles (e.g. late time points for N571) grew in X, Y and Z, creating high contrast phase images for which skeletonization revealed gaps between assemblies rather than assemblies themselves, necessitating image inversion. Typically, sample sets were compared by t-test in Excel. *p<0.05, **p<0.01, ***p<0.005.

### Immunofluorescence

Where fluorescence intensity is quantified, cells were imaged as a set with all images for each filter set captured with the same exposure time. Images were saved at 1499 by 1205 pixels in tagged image file format for downstream analysis. Image quantification was done on unmodified images; for area measures, region masks were generated in blue (nucleus) and green (CIZ1) using FIJI and area occupied by CIZ1 territories expressed as proportion of nuclear area. For frequency scores, typically analysis was carried out directly on samples, in triplicate, in two or more independent experiments with independent primary cell populations. For reproduction purposes, images were enhanced, split or cut using FIJI, in all cases to accurately reflect actual relationships between factors.

## RESOURCES

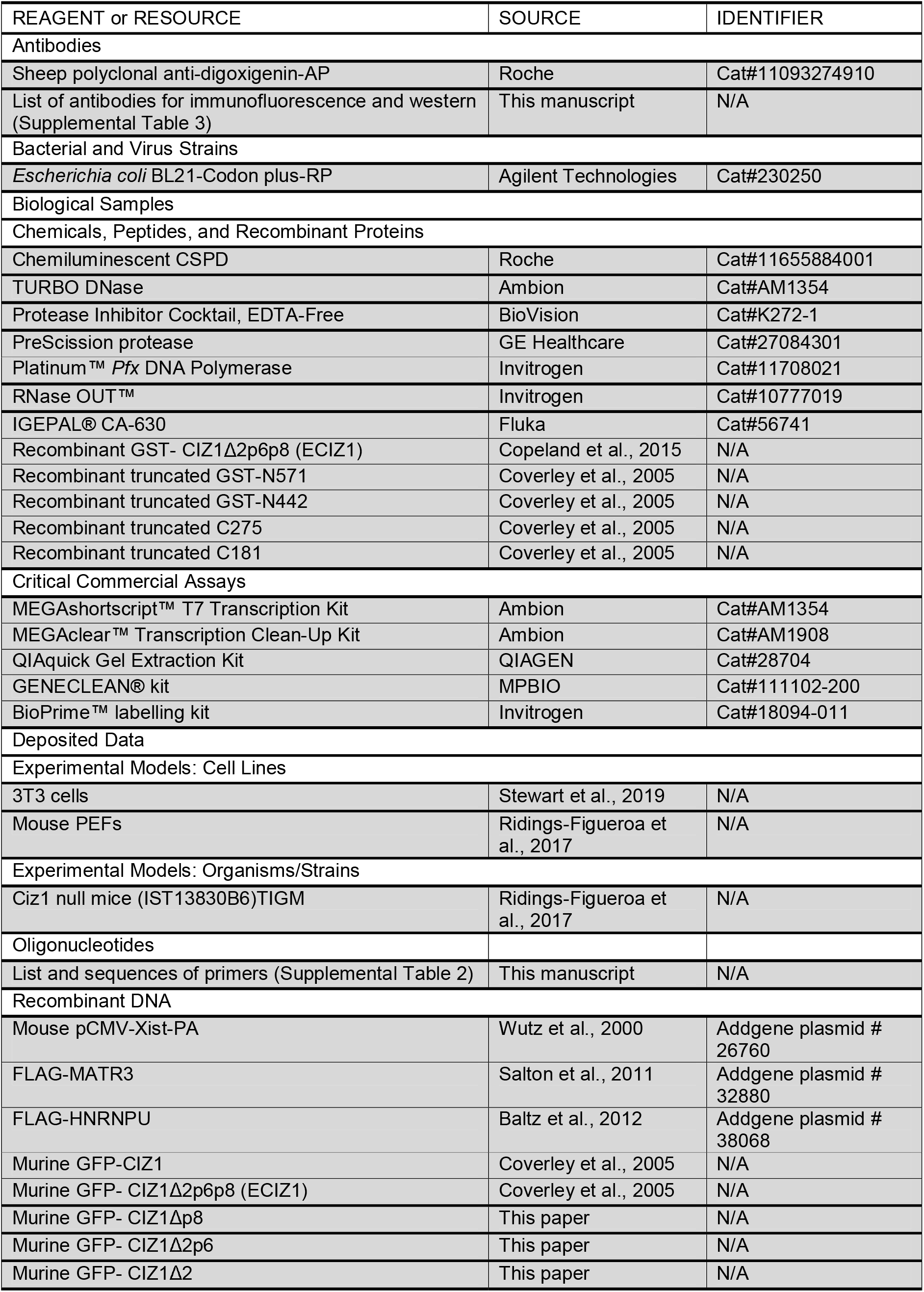
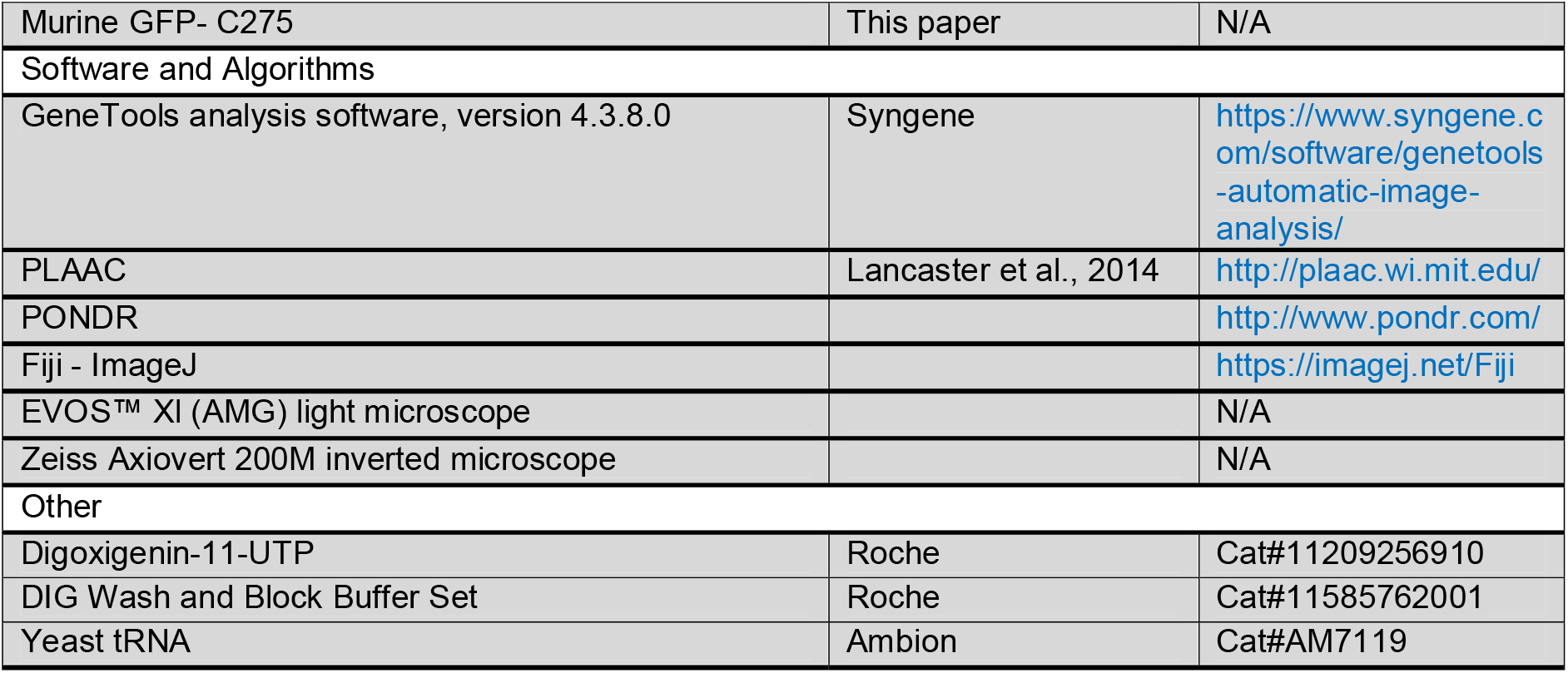

