## Supplemental text and figures for "A polyglutamine domain is required for *de novo* CIZ1 assembly formation at the inactive X chromosome"

#### Supplemental Information

##### Supplemental Fig. 1 (related to Fig. 1)

##### Sequence requirements for accumulation of murine GFP-CIZ1 at Xi

A) Quantification of the ability of the indicated constructs to form assemblies at Xi, after transient transfection into WT PEFs (p3), shown relative to H3K27me3-marked Xis which are evident in 80% of the total population. Results show mean frequencies from at least four experiments (N-values on images in B, with SEM and significance indicators calculated by t-test. B) Example images, acquired under standardised imaging conditions, showing H3K27me3-marked Xis (red), and GFP-CIZ1 assemblies (green) which, for the variant forms, are not only reduced in frequency, but when present are typically smaller, more diffuse entities (example nucleus with weak assembly is shown for CIZ1$\Delta$2). Values in A include weak assemblies. C) Representative images (related to main Figure 1D which gives mean % values, ±SEM) showing the behavior of the indicated GFP-CIZ1 variants (green) counterstained with DAPI (blue), 24 hours after transient transfection into 3T3 cells. Bar is 10 microns. Arrows indicate Xis with accumulated GFP. All constructs, except GFP empty vector, are exclusively nuclear and resistant to pre-fixation detergent exposure. Image capture parameters were not standardized across the different constructs.

**Supplemental Fig. 2 (related to Fig. 1)**

**Validation of nuclear import signal**

A) Exon map of human CIZ1 showing location of two putative nuclear localization signals (NLS, starred) detected by PSORTII; a classical NLS in the replication domain of CIZ1 (RD-NLS) encoded by exon 7, and a bipartite NLS (Robbins et al., 1991) in the anchor domain of CIZ1 (AD-NLS), encoded by exon 13. Above, sequences encompassing the predicted NLS (amino-acid positions from human protein reference sequence BAA85783.1), with key amino-acids in red. Below, products of mutagenesis with changed amino-acids in red. B) Proportion of transfected cells in which nuclear GFP fluorescence exceeds cytoplasmic fluorescence, for the indicated four constructs, 24 hours after transient transfection into male NIH3T3 cells (mouse) or female MCF7 cells (human). N and C refer to the individually mutated NLS's, double refers to mutation of both in the same construct. C) Representative images showing GFP-CIZ1 (green) without and with N-terminal NLS mutation, in NIH3T3 cells. Bar is 10 microns. D) Alignments showing human and mouse CIZ1 exon 7 and exon 13, with conserved NLS's boxed in red.

**Supplemental Fig. 3 (related to Fig. 1)**

**Similarity between mouse and human CIZ1**

A) Schematic comparing mouse and human CIZ1 exon structure, and validated excluded exons 2 (Coverley et al., 2005; Warder and Keherly, 2003), 4 (Greaves et al., 2012; Rahman et al., 2007; Warder and Keherly, 2003), 8 (Coverley et al., 2005; Dahmcke et al., 2008), 6 (Coverley et al., 2005; Greaves et al., 2012; Xiao et al., 2012) and 14 (Higgins et al., 2012) in green, or red for those which overlap with PLDs. PLD sequences align with alternatively spliced exons 2 (PLD1) and 8 (PLD2) in both mouse and human CIZ1. Predicted Atrophin 1 homology domain (cl26464) at amino-acids 121-482 in human CIZ1 (recently reclassified as cl33720**)** is shown by grey line. Exon 1 is untranslated and represented by at least three alternative versions in both human and mouse, and not shown here. B) Prion-like domains (PLDs) identified by PLAAC (Lancaster et al., 2014) are similar in human and mouse CIZ1. C) Disorder prediction by PONDR, showing overall 68.05% structural disorder for murine CIZ1 and 69.38% for human CIZ1. D) Predicted full-length murine (RefSeq: NP_082688.1) and human CIZ1 (BAA85783.1) amino-acid sequence, showing glutamine rich PLD1 and 2 in red, and conditionally excluded exons 2 and 8 in bold.

**Supplemental Fig. 4 (related to Fig. 3)**

**Supporting EMSA data**

A) Map showing mouse *Xist* (NR_001463.3) and 1.5 kb long *Xist* repeat E region (highlighted in blue). Repeat E regions implicated in CIZ1 recruitment as used by Sunwoo et al 2017 (red) and Ridings-Figueroa et al 2017 (light blue) are indicated. Repeat E probes proximal (dark blue) and distal (grey) used in this paper to test for direct CIZ1 binding are indicated below. B) Schematic representation of domain architecture of mouse CIZ1 and the truncated versions used here. PLD1 and PLD2 in mouse CIZ1 (NP_082688.1) represent prion like domains. ZnF1-3 zinc finger domain 1, zinc finger domain 2 and zinc finger domain 3. ZnF3 is also annotated as Jasmonate ZIM-domain (JAZ) by NCBI's Conserved Domain Database (CDD) https://www.ncbi.nlm.nih.gov/Structure/cdd/wrpsb.cgi. AcD represents acidic domain and Matrin matrin 3 domain. C) No binding was observed between C275 and *Gapdh* or 18S rRNA by EMSA. D) CIZ1Δ2p6p8 and C275 protein fragments bind proximal repeat E RNA as shown by EMSA. RNA-binding is lost upon heating to 95°C for 3 min prior to sample loading. Addition of 10 mM EDTA to binding reactions enhances binding. 6.5 nM RNA probe was used in all lanes. E, F) Truncated C-terminal fragment C181 interaction with *Xist* repeat A and repeat E (distal sense) probes respectively, and G) lack of interaction with *Gapdh* (analysis as for C275 in Fig. 3). H) EMSA blot used as an example to show which RNA fractions (red rectangles) were used to generate graphs for Figure 3 B,C,D. I) EMSA showing GST-N571 binding with *Gapdh*, but not 18S rRNA. RNA concentration used was 0.3 nM. All experiments were performed three times, carried out on different days.

**Supplemental Fig. 5 (related to Fig. 4)**

**Supporting *in vitro* assembly data**

A) Example phase contrast images of N571 (left) and N442 (right) at the indicated times and concentrations, showing emergence of mesh-like assemblies for N571, but not for N442. Bar is 10 microns. B) Histograms show the mean size, circularity and number of particles formed in samples of N571, incubated at the indicated concentrations for 24 hours. Error bars show SEM from 5 samplings. C) N571 incubated at 10 µM for 24 hours with and without tRNA at 500 ng/µl. Images were skeletonised as in Fig. 4C to extract information on size and the length of branches. Individual outputs for five samplings per condition are shown, compared by t-test. D) Control protein (bovine serum albumin) and RNA (total cellular RNA from female cultured fibroblasts) imaged after 24 hours at the indicated concentrations.

**Supplemental Fig. 6 (related to Fig. 5)**

***Xist*-interactors are not enriched in CIZ1 assembly zones**

A) Schematic representation of sequence elements in *Xist* interacting proteins PTBP1 (NP_032982.2), SAF-A, (NP_058085.2) and FLAG-MATR3 (NP_034901.2) showing PLDs (dark red), B) Left, immunofluorescence detection of CIZ1, SAF-A or PTBP1 (green) with co-staining for H3K27me3 (red) to mark Xi chromatin, in WT PEFs (p3), treated with detergent prior to fixation. Right, high magnification view showing lack of enrichment of SAF-A (green), and PTBP1 (green) in cells co-stained with CIZ1 (red). Arrows in detail images indicate CIZ1 assembly zones. DNA is blue, bar is 5 microns. C) Expression of ectopic Flag tagged MATR3 (green) in female 3T3 cells co-stained for endogenous CIZ1 (red). DNA is blue. Bar is 5 microns.

**Supplemental Table 1**

Summary of reported pathology-associated sequence variations in human CIZ1 PLD domains, including those documented more than once in COSMIC human tumour samples (Tate et al., 2019).

| Sequence variation | Disease | source |
|---|---|---|
| In frame deletion of Q9 in PLD1 | Adrenal gland Malignant Pheochromocytoma<br>Large intestine Adenocarcinoma<br>Large intestine Adenocarcinoma<br>Liver Carcinoma<br>Malignant melanoma<br>Malignant melanoma<br>Brain Haemangioblastoma | GRCh38·COSMIC v90 |
| SNP in PLD1 (exon 2) generating L36P | Thyroid Carcinoma | GRCh38·COSMIC v90 |
| SNP in PLD1 (exon 2) generating P47S | Cervical dystonia | (Xiao et al., 2012) |
| SNP in PLD1 (exon 2) generating R57W | Tongue Squamous cell carcinoma<br>Tongue Squamous cell carcinoma | GRCh38·COSMIC v90 |
| SNP in PLD1 (exon 2) generating L59P | Astrocytoma Grade IV<br>Astrocytoma Grade IV<br>Glioma<br>renal cell carcinoma | GRCh38·COSMIC v90 |
| Exclusion of PLD 1 (exons 2, 4, 6) | Medulloblastoma | (Warder and Keherly, 2003) |
| SNP in (exon 7) S264G with effects on splicing and nuclear aggregate formation | Cervical dystonia | (Xiao et al., 2012) |
| SNP in PLD2 (exon 8) generating Q360* | Lung Adenocarcinoma<br>Stomach Adenocarcinoma | GRCh38·COSMIC v90 |
| Conditional exclusion of part of PLD2 (part exon 8) | Alzheimer's disease | (Dahmcke et al., 2008) |

**Supplemental Table 2**

Primers used to generate DNA templates for *in vitro* transcription.

| probe | length | forward | reverse |
|---|---|---|---|
| RepA sense | 1-500 | T7-F: TAATACGACTCACTATAGGGAGCTTGCTCCAGCCATGTTTGCTCG | R: CTAAGGAGAAGAAAAAAAGAATAAAAGC |
| RepE proximal | 1-474 | T7-F: TAATACGACTCACTATAGGGAGATTTCTTCCTTGCAGTTGTGTCTAATTC | R: ACAGAGAGCCATAGCTAGTGAAG |
| RepE distal sense (20 nucleotide overlap with RepE proximal) | 1-472 | T7-F: TAATACGACTCACTATAGGGAGCACTAGCTATGGCTCTCTG | R: CACATAACACACATGCACACACGC |
| RepE distal antisense | 1-472 | F: CACTAGCTATGGCTCTCTGTTTTATCTATCTG | T7-R: TAATACGACTCACTATAGGGAGCACATAACACACATGCACACACGC |

**Supplemental Table 3**

Antibodies used for immunofluorescence and western blot studies.

| **Antibodies** | **Source** |
|---|---|
| 1794 (N-term CIZ1) | Coverley et al., 2005 |
| C- term mCIZ1 | Novus (NB100-74624) |
| C- term hC221a | (Stewart et al., 2019) |
| H3K27me3 | Abcam (6002) |
| H3K27me3 | Abcam (9733S) |
| H2AK119UB | CST (8240) |
| FLAG-tag antibody | Biorbyt (orb323042) |
| SAFA | Abcam ab10297 |
| lamin B | Invitrogen 33-2100 |
| anti-GST | Abcam ab9085 |
| Goat α Rabbit Alexa Fluor 568 (Red) | Invitrogen (A11011) |
| Goat α Rabbit Alexa Fluor 488 (Green) | Invitrogen (A11034) |
| Goat α Mouse Alexa Fluor 568 (Red) | Invitrogen (A11031) |
| Goat α Mouse Alexa Fluor 488 (Green) | Invitrogen (A11001) |
| Peroxidase IgG α Rabbit | Jackson 211-032-171 |
| Peroxidase IgG α Mouse | Jackson 155-035-174 |

### Supplemental Fig. 1

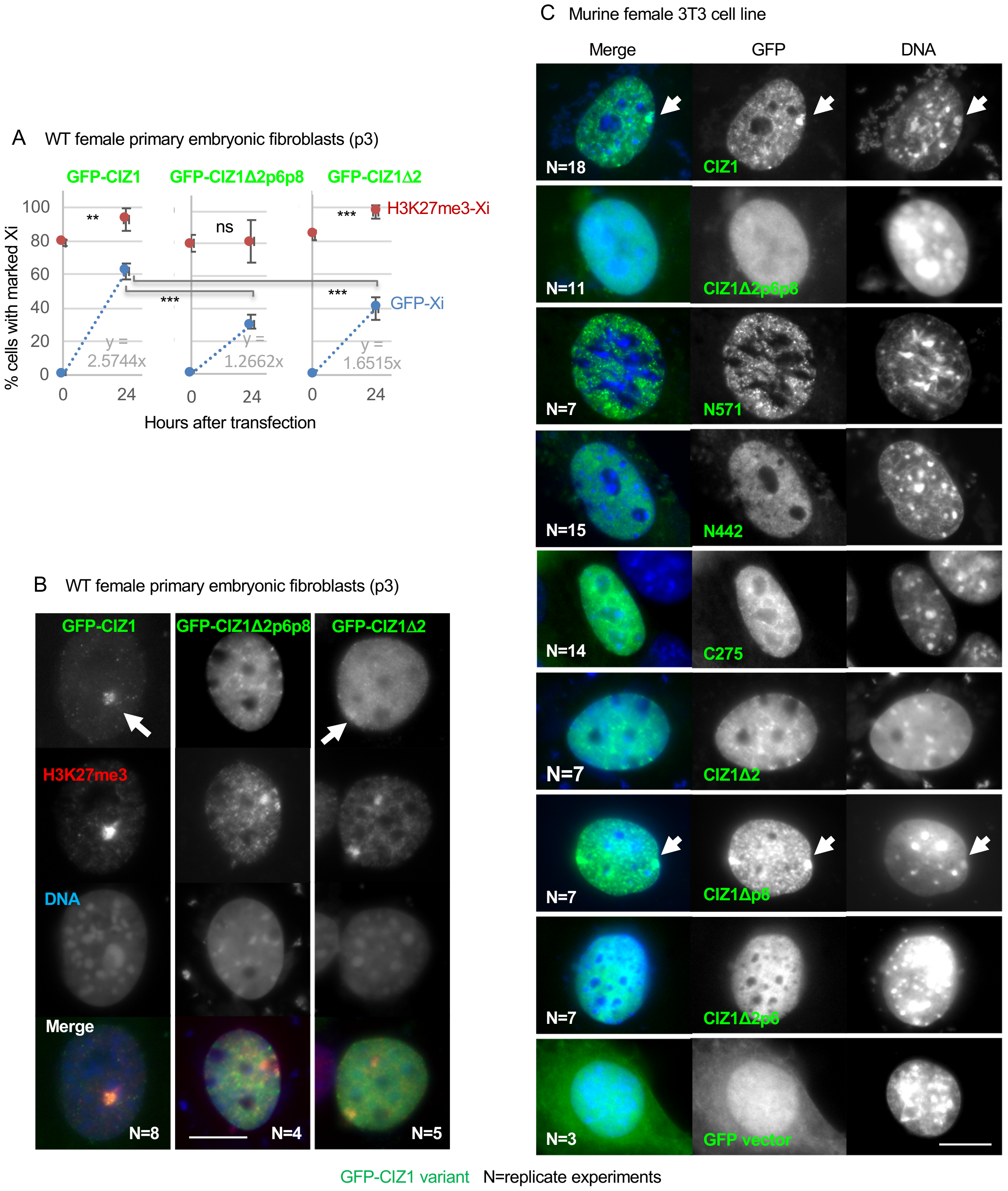

### Supplemental Fig. 2

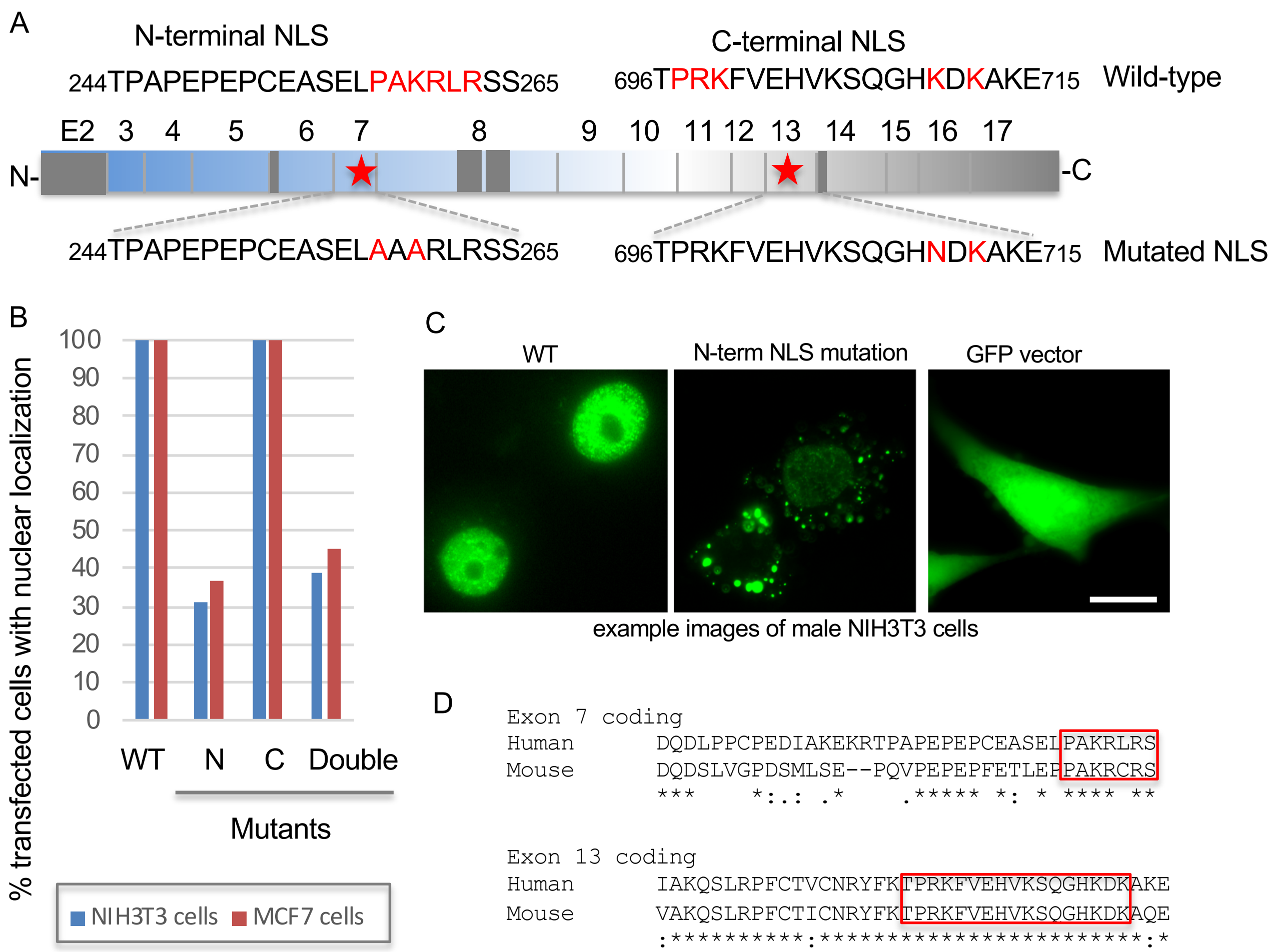

### Supplemental Fig. 3

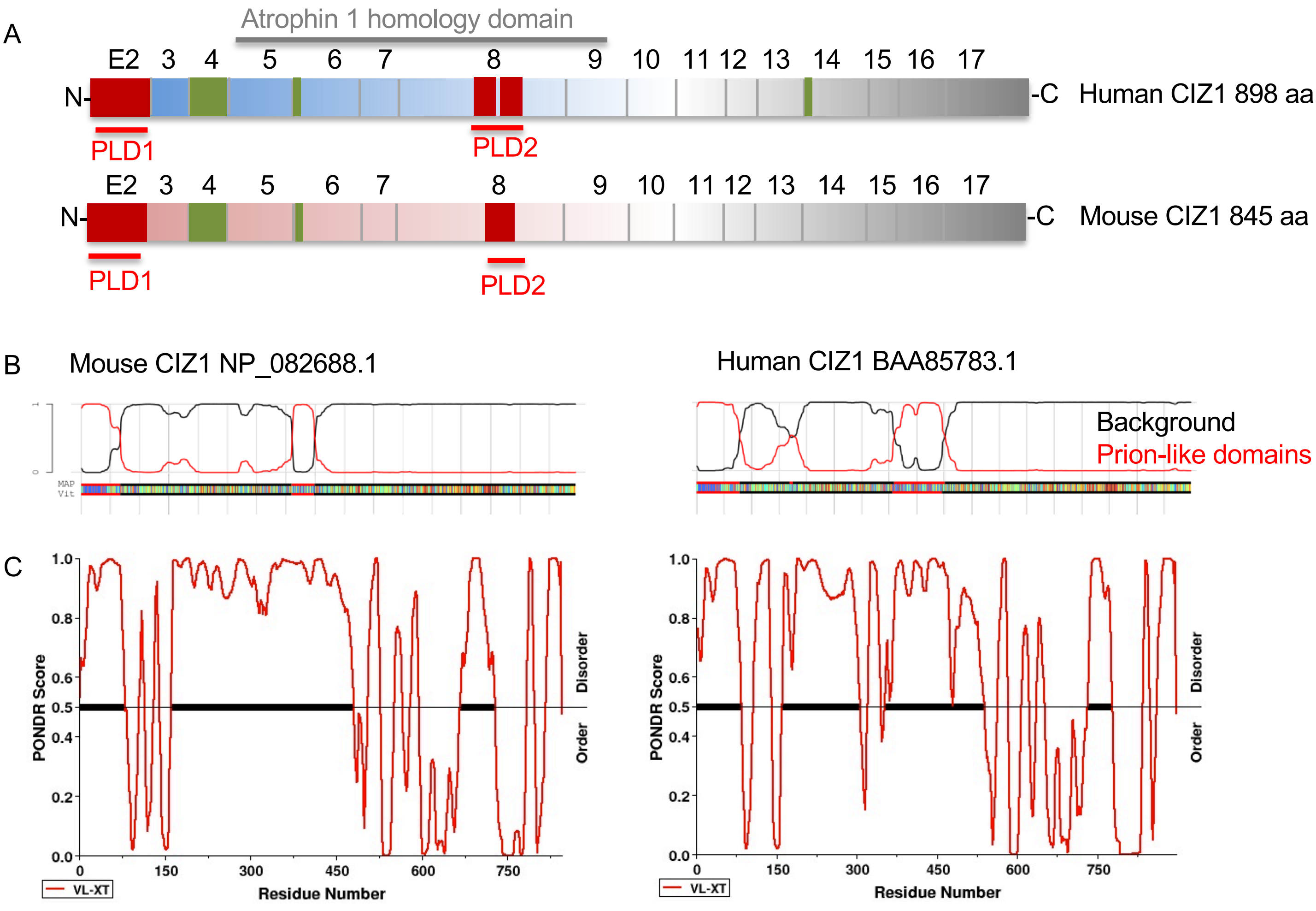


D. Mouse CIZ1 NP_082688.1, Human CIZ1 BAA85783.1

```
Mouse CIZ1   MFNPQLQQQQQLQQQQQQLQQ--QLQQQQLQQQQQQILQLQQLLQQSPPQASLSIPVSRGLPQQSSPQQLLSLQGLHSTSLLNGPMLQRA
Human CIZ1   MFSQQQQQ--QLQQQQQQLQQLQQQQLQQQQLQQQQLLQLQQLLQQSPPQAPLPMAVSRGLPPQQPQQPLLNLQGTNSASLLNGSMLQRA

Mouse CIZ1   LLLQQLQGLDQFAMPPATYDGASLTMPTATLGNLRAFNVTAPSLAAPSLTPPQMVTPNLQQFFPQATRQSLLGPPPVGVPINPSQLNHSG
Human CIZ1   LLLQQLQGLDQFAMPPATYDTAGLTMPTATLGNLRGYGMASPGLAAPSLTPPQLATPNLQQFFPQATRQSLLGPPPVGVPMNPSQFNLSG

Mouse CIZ1   RNTQKQARTPSSTTPNRKDSSSQTVPLEDREDPTEGSEEATELQMDTCEDQDSLVGPDSMLSEPQ--VPEPEPFETLEPPAKRCRSSEES
Human CIZ1   RNPQKQARTSSSTTPNRKDSSSQTMPVEDKSDPPEGSEEAAEPRMDTPEDQDLPPCPEDIAKEKRTPAPEPEPCEASELPAKRLRSSEEP

Mouse CIZ1   TEKGPTGQPQARVQPQTQMTAPKQTQTPDRLPEPPEVQMLPRIQPQALQIQ-----------------TQPKLLRQAQTQTSPEHLAPQQ
Human CIZ1   TEKEPPGQLQVKAQPQARMTVPKQTQTPDLLPEALEAQVLPRFQPRVLQVQAQVQSQTQPRIPSTDTQVQPKLQKQAQTQTSPEHLVLQQ

Mouse CIZ1   DQVEPQVPSQPPW--QLQPRE-----TDPPNQAQAQTQPQPLW------------------QAQSQKQAQTQAHPQVPTQAQSQ------
Human CIZ1   KQVQPQLQQEAEPQKQVQPQVQPQAHSQGPRQVQLQQEAEPLKQVQPQVQPQAHSQPPRQVQLQLQKQVQTQTYPQVHTQAQPSVQPQEH

Mouse CIZ1   ------EQTSEKTQDQPQTWPQGSVPPPEQASGPACATEPQLSSHAAEAGSDPDKALPEPVSAQSSEDRSREASAGGLDLGECEKRAGEM
Human CIZ1   PPAQVSVQPPEQTHEQPHTQPQVSLLAPEQTPVVVHVCGLEMPPDAVEAGGGMEKTLPEPVGTQVSMEEIQNESACGLDVGECENRAREM

Mouse CIZ1   LGMWGAGSSLKVTILQSSNSRAFNTTPLTSGPRPGDSTSATPAIASTPSKQSLQFFCYICKASSSSQQEFQDHMSEAQHQQRLGEIQHSS
Human CIZ1   PGVWGAGGSLKVTILQSSDSRAFSTVPLTPVPRPSDSVSSTPAATSTPSKQALQFFCYICKASCSSQQEFQDHMSEPQHQQRLGEIQHMS

Mouse CIZ1   QTCLLSLLPMPRDILEKEAEDPPPKRWCNTCQVYYVGDLIQHRRTQEHKVAKQSLRPFCTICNRYFKTPRKFVEHVKSQGHKDKAQELKT
Human CIZ1   QACLLSLLPVPRDVLETEDEEPPPRRWCNTCQLYYMGDLIQHRRTQDHKIAKQSLRPFCTVCNRYFKTPRKFVEHVKSQGHKDKAKELKS

Mouse CIZ1   LEKETGSPDEDHFITVDAVGCFESGQEEDEDDDEEEEEEGEIEAEEEFCKQVKPRETSSEQGKGSETYNPNTAYGEDFLVPVMGYVCQIC
Human CIZ1   LEKEIAGQDEDHFITVDAVGCFEGDEEEEEDDE----DEEEIEVEEELCKQVRSRDISREEWKGSETYSPNTAYGVDFLVPVMGYICRIC

Mouse CIZ1   HKFYDSNSELRLSHCKSLAHFENLQKYKAKNPSPPPTRPVSRKCAINARNALTALFTSSHQPSPQDTV--KMPSKVKPGSPGLPPPLRRS
Human CIZ1   HKFYHSNSGAQLSHCKSLGHFENLQKYKAAKNPSPTTRPVSRRCAINARNALTALFTSSGRPPSQPNTQDKTPSKV-TARPSQPPLPRRS
Mouse CIZ1   TRLKT
Human CIZ1   TRLKT
```

### Supplemental Fig. 4

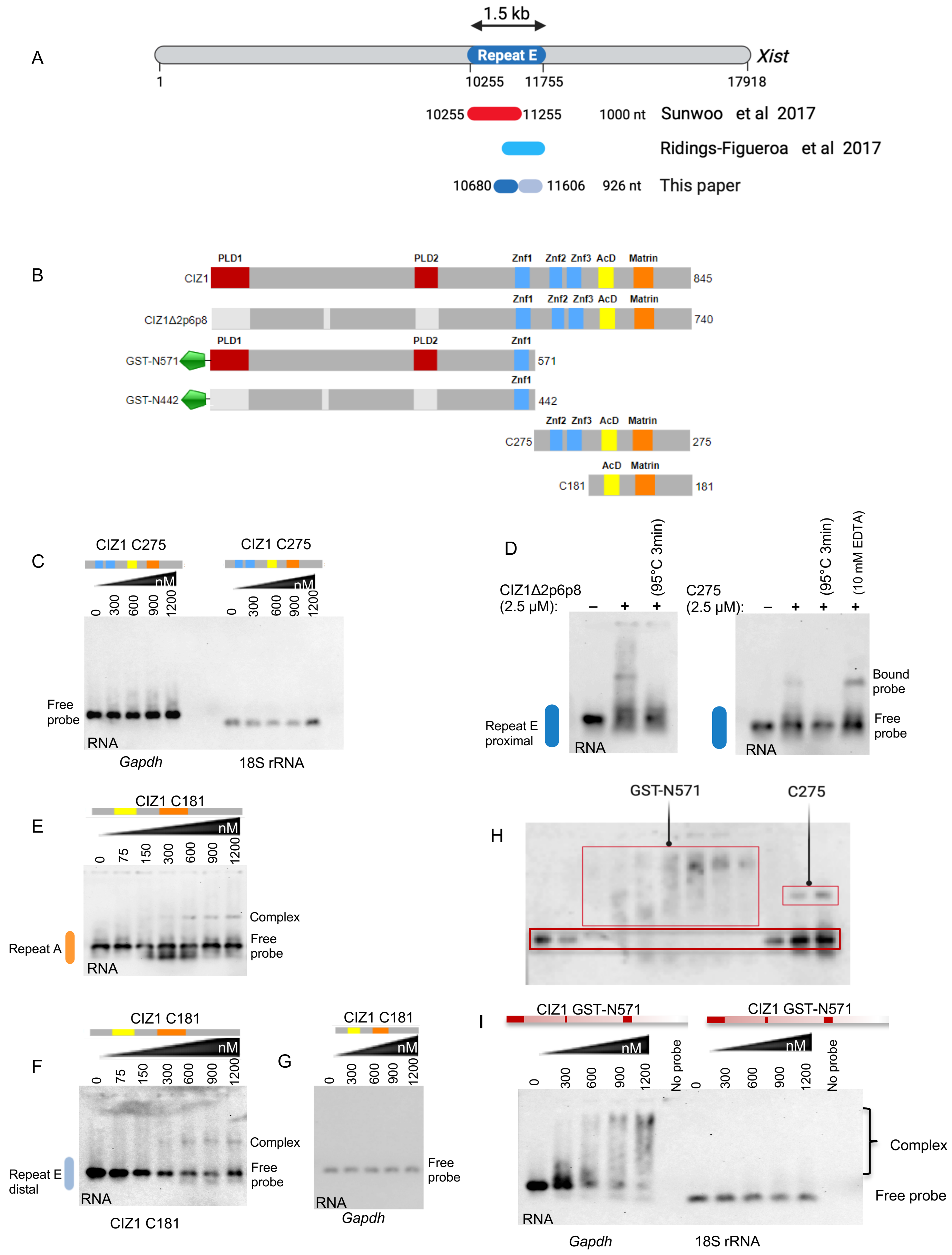

### Supplemental Fig. 5

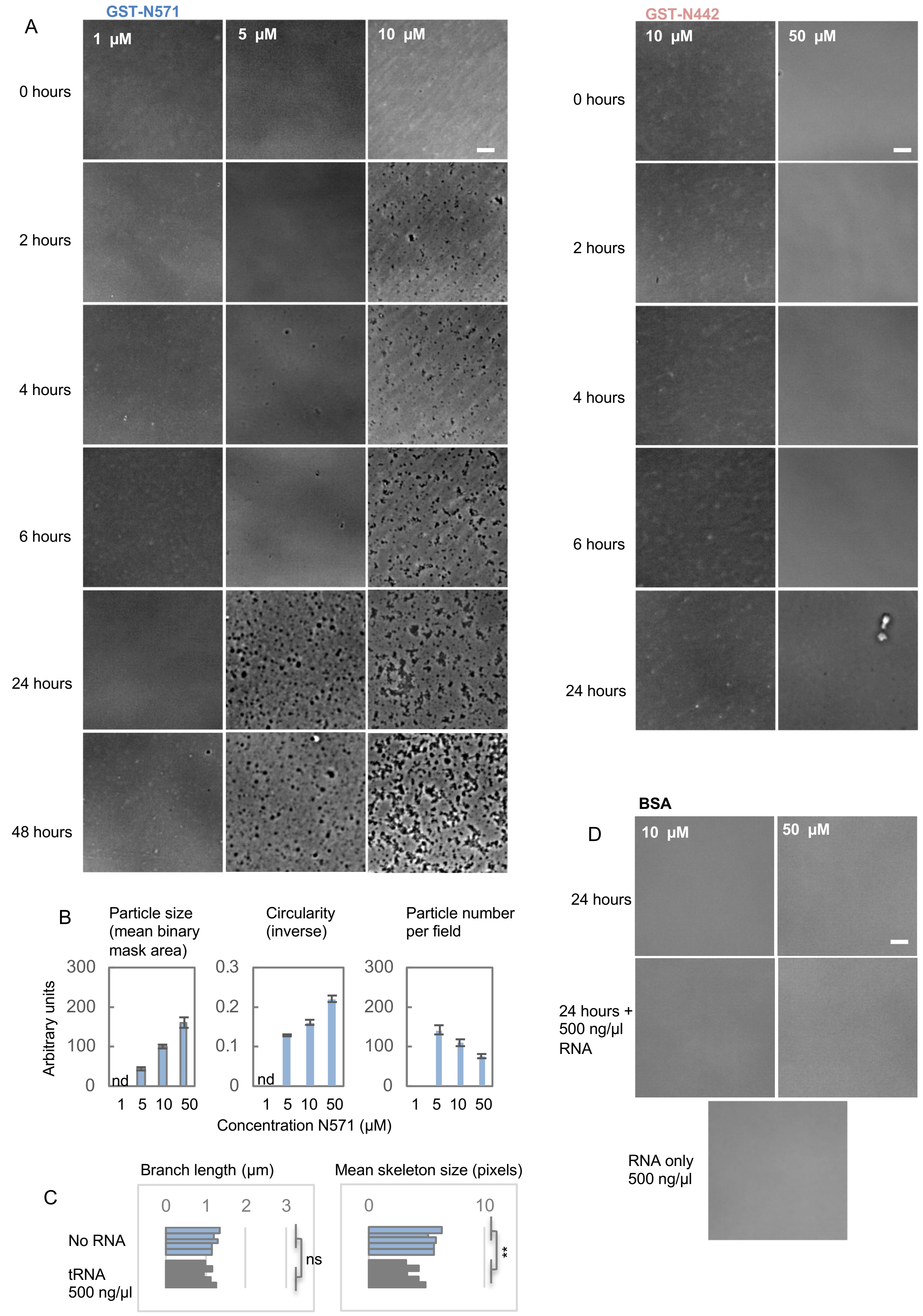

### Supplemental Fig. 6

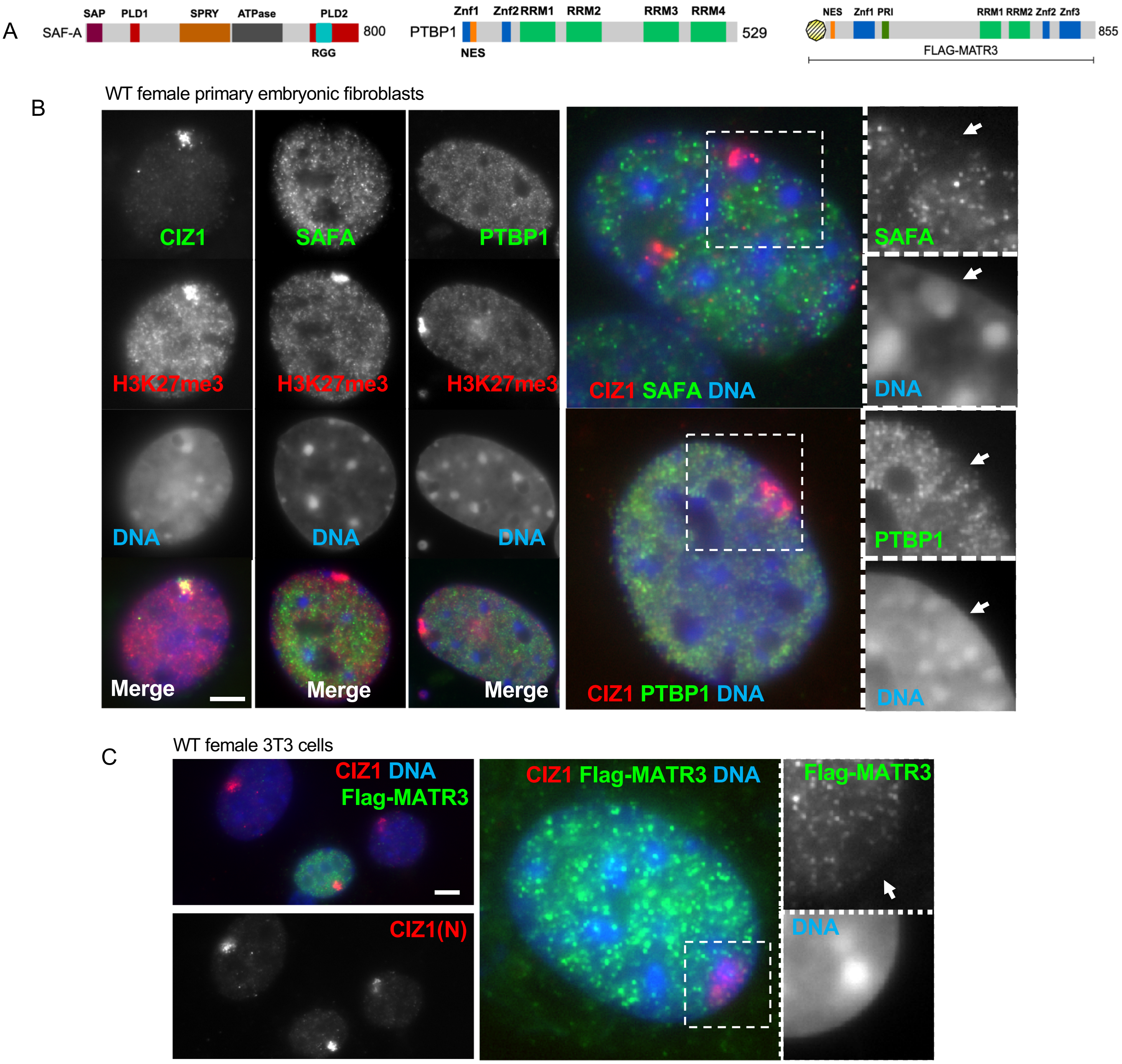
